# Diversity of penicillin-binding protein 2x in *Streptococcus pyogenes* from England and Wales, 2001 to 2015

**DOI:** 10.64898/2026.09.21.752073

**Authors:** Greta Šveikauskaitė, Asha L Marshall, Sofya Gorshkova, Rebecca L Guy, Yu Wan, Ana Vieira, Juliana Coelho, Kartyk Moganeradj, Shiranee Sriskandan, Elita Jauneikaite, Valerie WC Soo

**Affiliations:** Department of Infectious Disease, Imperial College London, UK; Centre for Bacterial Resistance Biology, Imperial College London, UK; NIHR Health Protection Research Unit (NIHR HPRU) in Healthcare Associated Infections and Antimicrobial Resistance, Department of Infectious Disease, Imperial College London, UK; Antimicrobial Resistance and Healthcare Associated Infections Division, UK Health Security Agency, UK; David Price Evans Global Health and Infectious Diseases Research Group, University of Liverpool, UK; Staphylococcus & Streptococcus Reference Section, UK Health Security Agency, UK; Department of Metabolism, Digestion and Reproduction, Imperial College London, UK

**Keywords:** Group A *Streptococcus*, penicillin-binding protein, antibiotic target, antimicrobial resistance, β-lactam antibiotics, multidrug resistance

## Abstract

*Streptococcus pyogenes* remains clinically susceptible to β-lactam antibiotics, its first-line treatment. Yet recent reports from the US, Iceland, and Japan have identified isolates with PBP2x (a primary β-lactam target) substitutions that reduced antibiotic susceptibility. To determine whether these PBP2x substitutions occurred in *S. pyogenes* from England and Wales, we re-analysed 2,970 invasive genomes from 2001 to 2015. We found 42 PBP2x sequence mutants in 34% (996/2,970) of the genomes, revealing an unexpected diversity in this antibiotic target. Nine *emm*12 isolates carried a novel double substitution, PBP2x G600A_P601H, that co-occurred with macrolide resistance genes. Experimentally, these isolates showed 2- to 4-fold elevated minimum inhibitory concentrations for four β-lactam antibiotics, alongside erythromycin resistance. Other PBP2x substitutions, found across seven *emm* types without reduced antibiotic susceptibility, appeared driven by *emm* lineage expansion rather than antibiotic pressure. These findings establish a baseline for *pbp*2x variation, reinforcing the case for sustained genomic surveillance globally.

## INTRODUCTION

*Streptococcus pyogenes*, also commonly known as Group A *Streptococcus*, has evolved to exclusively colonise and infect humans. Within this single host, *S. pyogenes* can thrive in many body tissues, as exemplified by its diverse range of disease manifestations^1^. Mild infections such as pharyngitis, impetigo, and scarlet fever are self-limiting and treatable with antibiotics. Severe, invasive infections such as necrotizing fasciitis, myositis, streptococcal toxic shock syndrome, and sepsis have an average mortality rate of approximately 20%^2,3^. Further, recurrent episodes of *S. pyogenes* infections or unexpected clinical complications can trigger the onset of immune-mediated post-infectious sequelae, including acute rheumatic fever, rheumatic heart disease, and post-streptococcal glomerulonephritis^1^. Such a broad spectrum of diseases places a significant burden on public health worldwide. Coupled with cyclical and frequent upsurges of *S. pyogenes* infections since the mid-2010s^4,5^, *S. pyogenes* is now one of the 15 most significant bacterial pathogens contributing to global number of deaths and years of life lost^6^.

Without a safe and effective vaccine^7^, antibiotics remain the only effective therapeutic option for *S. pyogenes* infections. Penicillins, including phenoxymethylpenicillin, benzylpenicillin, and amoxicillin (all are β-lactam antibiotics), are the first-line antibiotics used for treating *S. pyogenes* infections in all countries. Second-line antibiotics (such as macrolides) are considered for patients with severe penicillin allergy. Clindamycin, a lincosamide antibiotic, is occasionally used as an adjuvant therapy along with penicillins. However, antimicrobial resistance is an emerging problem for *S. pyogenes* in many countries, which resulted in its inclusion as a priority pathogen by the World Health Organization^8^ and the UK Health Security Agency^9^. In the UK, laboratory surveillance data from 2024/2025 indicated that 18% of the invasive isolates of *S. pyogenes* were resistant to erythromycin, 8% were resistant to clindamycin, and 41% were resistant to tetracyclines^10^. Similarly, invasive *S. pyogenes* isolates from the Active Bacterial Core surveillance in the US have been showing increasing rates of erythromycin resistance and clindamycin resistance since mid-2015^11,12^. Cross resistance is particularly evident in China, where up to 97% of *S. pyogenes* isolates were resistant to both erythromycin and clindamycin in multiple provinces^13^. Whilst all *S. pyogenes* isolates remain clinically susceptible to β-lactam antibiotics, reduced susceptibility to this antibiotic class is a major clinical and public health concern. Treatment failures with β-lactam antibiotics are not uncommon, even for non-invasive diseases^14^.

The development of β-lactam resistance in other streptococcal pathogens requires an initial acquisition of non-synonymous mutations in the essential *pbp2x* gene, which encodes for the bacterial penicillin-binding protein 2x (PBP2x)^15,16^. As a primary inhibition target of β-lactam antibiotics, PBP2x is a transpeptidase enzyme that crosslinks peptidoglycan strands during bacterial cell wall biosynthesis. In *Streptococcus pneumoniae*, non-synonymous mutations resulting in altered PBP2x may lead to a reduced binding^17^, or an increased turnover^18^, of a β-lactam antibiotic. Hence, it is important to investigate whether any of the PBP2x substitutions may represent a feasible mutational route to β-lactam non-susceptibility in *S. pyogenes*.

Genomic and epidemiological studies from the US^19–22^, Iceland^23^, and Japan^24^ have reported *S. pyogenes* isolates with a subset of *pbp2x* mutations showing reduced levels of susceptibility to β-lactam antibiotics. Two *S. pyogenes* isolates expressing PBP2x with a T553K substitution exhibited an 8-fold increase in minimum inhibitory concentrations (MICs) for amoxicillin and ampicillin as well as a 3-fold increase for cefotaxime^19^. Other substitutions showing minor but consistent increases in MICs to various β-lactam antibiotics (1.5- to 2-fold) include P601L^22,25,26^, M593T^20,22–24,26^, G600D^25,26^, and A397V^24^. Despite these elevated MICs, all isolates have remained below the clinical resistance breakpoints for *S. pyogenes*, as defined by the CLSI^27^ and EUCAST guidelines^28^.

In the UK, antimicrobial susceptibility is routinely reported for all invasive clinical isolates of *S. pyogenes*^10^; however subtle changes in β-lactam antibiotic susceptibility would not be detected and the frequency of PBP2x substitutions unknown. In this study, we analysed two published national collections of invasive *S. pyogenes* genomes spanning 2001 to 2015 from England and Wales. Focusing on the prevalence of non-synonymous mutations in *pbp2x*, we captured a moderate level of PBP2x diversity in this dataset, with select isolates showing elevated MICs to various β-lactam antibiotics. Some PBP2x variants were also associated almost exclusively with certain *emm* types, highlighting a low risk of *pbp2x* diversification along with future expansion of certain *emm* lineages.

## RESULTS

### Diversity of invasive S. pyogenes in England and Wales from 2001 to 2015

All cases of scarlet fever and invasive Group A Streptococcal (iGAS) disease in the UK are notifiable under the Health Protection (Notification) Regulations 2010. Around the same time, whole-genome sequencing became common and has since been deployed as a public health measure, e.g., to track outbreaks of *S. pyogenes*, to identify the dominant or high-risk circulating lineages, to infer virulence potential, and to assess the transmission of AMR determinants.

To understand the prevalence and diversity of AMR determinants of *S. pyogenes* in the UK, we retrospectively analysed two published genome datasets consisting of 3,391 invasive isolates of *S. pyogenes* collected in England and Wales from 2001 to 2015. Sequencing reads from a total of 2,970 isolates [318 from the BSAC Bacteraemia Resistance Surveillance Program^29^ and 2,652 from the formerly known Public Health England National Streptococcal Reference Laboratory^30^] passed our quality checks and were assembled for molecular typing and comparative analyses. We identified 74 *emm* types, with *emm*1 (21.9%) being the most prevalent *emm* type (Supplementary Table 1, Figure 1a). This was followed by *emm*3 (17.7%), *emm*12 (12.4%), *emm*28 (7.4%), *emm*75 (4.8%), *emm*89 (4.4%), *emm*4 (4.4%), *emm*6 (3.9%), *emm*87 (2.7%), and *emm*5 (2.3%) (Supplementary Table 1, Figure 1A). These top 10 *emm* types accounted for 82% of the overall dataset, reflecting their prevalent circulation and outbreaks in England and Wales during the study period^29,30^. MLST revealed 149 Sequence Types (STs), predominantly ST28 (Supplementary Table 1, Figure 1B). Most STs and their single-locus variants (SLVs) and double-locus variants (DLVs) were associated with distinct *emm* types. For instance, each of the top ten *emm* types was dominated by a single ST (>86%), except for *emm*3, which was dominated by ST315 (53%) and one of its SLVs, ST15 (36%) (Figure 1C). Overall, these isolates were genomically diverse, capturing a moderate degree of strain variability (Figure 2).

**Figure 1.**
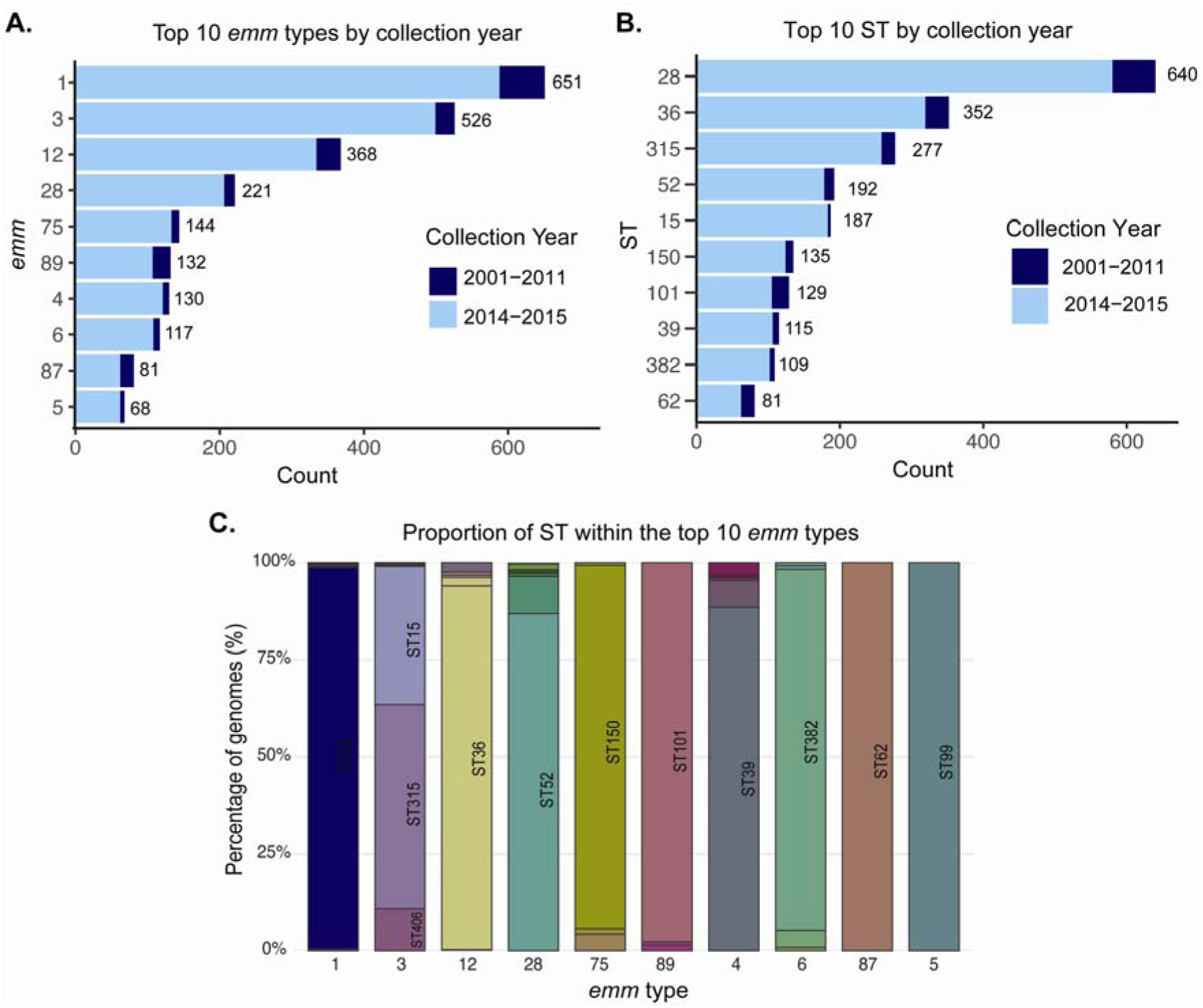
Variability of *emm* types and STs of all 2,970 invasive *S. pyogenes* genomes collected between 2001 and 2015 from England and Wales. (**A**) Top 10 *emm* types stratified by collection years. (**B**) Top 10 STs stratified by collection years. (**C**) Dominance of single STs within the top 10 *emm* types (except for *emm*3).

**Figure 2.**
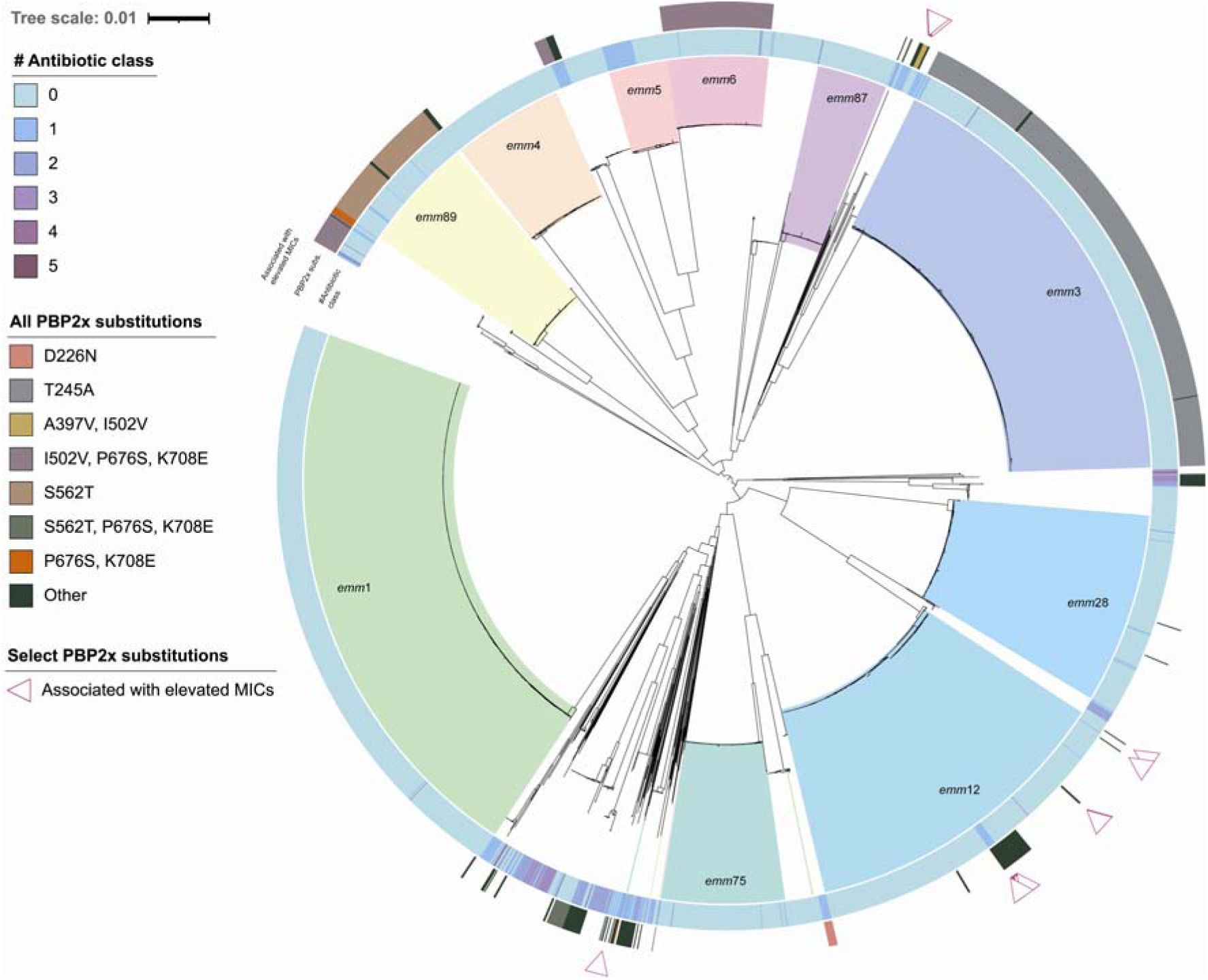
Maximum likelihood phylogenetic tree for 2,970 invasive *S. pyogenes* genomes collected between 2001 and 2015 from England and Wales. The tree was constructed from the alignment of 1,379 core genes generated by Panaroo. The majority of the *S. pyogenes* genomes clustered by *emm* types, and the top 10 *emm* types are highlighted and annotated accordingly. The inner ring denotes the number of antibiotic classes to which the *S. pyogenes* isolates were predicted to be resistant. PBP2x substitutions are represented by the middle ring, and those that are known to be associated with elevated MICs to β-lactam antibiotics are marked as pink triangular markers. The scale bar represents the mean number of nucleotide substitutions per site.

### Antibiotic resistance to macrolide, tetracycline, lincosamide, chloramphenicol, and trimethoprim/sulfamethoxazole

Within this dataset, 315 genomes (10.6% of 2,970 genomes) encoded 1-6 AMR genes (Inset, Figure 3) that could confer resistance to macrolides, tetracyclines, lincosamides, chloramphenicol, and trimethoprim/sulfamethoxazole. Phenotypic resistance to macrolides (erythromycin), lincosamides (clindamycin), and tetracycline was previously confirmed for a small subset of these isolates (55 isolates from the BSAC collection), in which phenotypic resistance was strongly correlated with detected AMR genes^29^. Among the 315 AMR gene-positive genomes, over half (n = 163; 52%) harboured a single AMR gene conferring resistance to either tetracyclines (n = 152) or macrolides (n = 11) (Figure 3). One genome belonging to *emm*81 carried six AMR genes (*tetM*, *ermB*, *aph3-III*, *ant6_Ia*, *cat_pC194*, and *lnuC*) which might result in resistance to five types of antibiotic classes (tetracyclines, macrolides, aminoglycosides, chloramphenicol, and lincosamides) (Figure 3).

**Figure 3.**
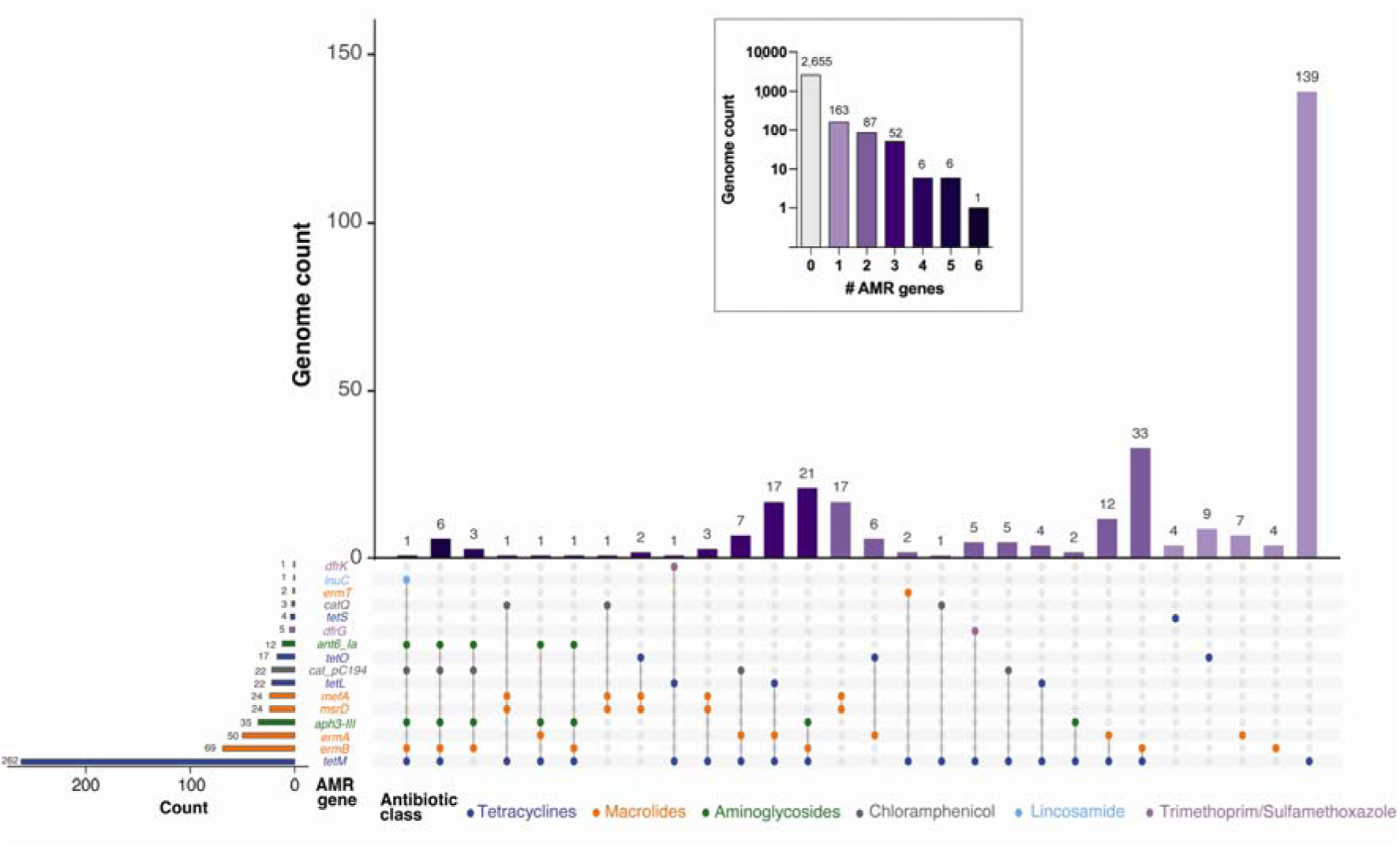
Distribution AMR gene profiles and their associated antibiotic class across the 315 AMR gene-positive *S. pyogenes* genomes. Inset: Number of AMR genes detected in all the *S. pyogenes* genomes (n = 2,970) in this study.

Although aminoglycosides and chloramphenicol are not used for any *S. pyogenes* infections, a small number of genomes carried AMR genes that could confer resistance to these antibiotic classes (Figure 3). Similarly, trimethoprim/sulfamethoxazole is not a routine antibiotic for *S. pyogenes*; however, its use is occasionally considered if the *S. pyogenes* isolate is erythromycin- and clindamycin-resistant and penicillin use is not possible under specific clinical circumstances. In our dataset, only six genomes encoded the *dfr* genes, which might confer resistance to trimethoprim/sulfamethoxazole (Figure 3). In this study, we observed 41 genomes that were predicted to be multidrug resistant, as their combinations of AMR genes could confer resistance to three or more antibiotic classes (Figure 3). Notably, 75 genomes were predicted to exhibit resistance to both tetracyclines and macrolides (Figure 3). Tetracycline resistance accounted for the highest number of AMR gene-positive genomes in our dataset. As second-line antibiotics and adjuvants, erythromycin and clindamycin resistance in *S. pyogenes* is high in China^13^ and the US^11^, but in this dataset we detected a small number of genomes having *ermB* (n = 69), *ermA* (n = 50) and *ermT* (n = 2) that could confer resistance to both macrolides and clindamycin (n = 121; 4.1% of the entire collection; Figure 3). One genome harboured the *lnuC* gene that confers resistance to lincosamides. Other macrolide resistance genes were *mefA* and *msrD* (n = 24).

To facilitate retrospective epidemiological studies, we analysed the breakdown of *emm* types for 315 AMR gene-positive genomes. We detected a total of 52 *emm* types with at least one AMR gene, accounting for 70% of all *emm* types in our dataset. Given the high prevalence of the top 10 *emm* types in this dataset (n = 2,438; 82%; Figure 1a), we expected a high proportion of AMR genes in these *emm* types accordingly. However, this was not observed; in fact, the top 10 *emm* types were responsible for only 24% (n = 76) of all AMR gene-positive genomes, implying that a majority of AMR genes were distributed across less common *emm* types in *S. pyogenes* (Pearson’s χ^2^_1df_ = 800.67, *P* = 3.86e-176; Figure 2). This finding was consistent with the rare occurrence of AMR determinants in the six most common *emm* types in the US^12^. Within the top 10 *emm* types, *emm*5 contained the highest proportion of AMR gene-positive genomes (n = 33/68; 48.5%; Supplementary Table 1, Figure 2). In contrast, no AMR gene was found in all *emm*4 and *emm*87 genomes. For the dominating *emm*1 lineages in the UK^31^, only 2 isolates (0.3% of 651 *emm*1 genomes) harboured AMR genes that conferred resistance to macrolides, tetracyclines, and aminoglycosides.

In the US, erythromycin resistance was driven by the expansion of the *emm*92 and *emm*83 lineages between 2016 and 2017^12^, with both types having been associated with socially vulnerable populations in the UK^32^. In our dataset, however, there were only three *emm*92 and three *emm*83 genomes, making it difficult to compare with the US finding. None of our *emm*92 and *emm*83 genomes harboured any macrolide resistance gene. Of the 145 genomes with macrolide resistance genes, the most prevalent *emm* types were *emm*11 (n = 39; 26.9%), *emm*168 (n = 17; 11.7%), and *emm*58 (n = 15; 10.3%). Similarly, most of the macrolide-resistant *S. pyogenes* from Europe during this period belonged to *emm*11 and *emm*58, both of which encoded macrolide resistance genes in integrative conjugative elements^33–36^.

### pbp2x mutations associated with reduced bacterial susceptibility to β-lactam antibiotics

Despite universal susceptibility to β-lactam antibiotics, *S. pyogenes* containing some non-synonymous mutations in *pbp2x* (T553K, M593T, P601L) can show, or have been associated with (A397V, P526S), elevated MIC values. In two *emm*43/ST3 isolates of *S. pyogenes*, a T553K substitution in one of the three conserved motifs of PBP2x (S_340_TMK, S_399_SN, and K_550_SGT) has led to the largest MIC increases in amoxicillin, ampicillin, and cefotaxime^19^. However, Hayes *et al.* noted that substitutions in any of these conserved motifs were rare^37^. Consistent with Hayes *et al.*, we also did not observe any non-synonymous mutations, including T553K, in these conserved motifs of PBP2x within our dataset. Furthermore, none of the 10 *emm*43/ST3 isolates within our dataset carried non-synonymous mutations in *pbp2x* (Supplementary Table 1). Mosaic *pbp2x* alleles commonly found in β-lactam-resistant strains of *S. pneumoniae*^38^ and *S. dysgalactiae*^39^ were also not present in this dataset.

Yet, in our dataset non-synonymous *pbp2x* mutations were present, and a moderate diversity in protein sequence types were observed. Excluding the ‘wildtype’ reference PBP2x sequence from the penicillin-susceptible *emm*1 MGAS5005, a total of 42 PBP2x sequence variants were detected in 33.5% genomes (n = 996) of our entire collection Supplementary Table 1; Supplementary Data). These sequence variants were encoded by 102 *pbp2x* alleles (Supplementary Table 1). The number of PBP2x substitution ranged from one to four, with substitutions found throughout the four protein domains of PBP2x [transmembrane, PBP_dimer (Pfam PF03717), transpeptidase (Pfam PF00905), and two PASTA (Pfam PF03793) domains] as well as interdomain linkers (Figures 4A). Among the protein domains, PBP_dimer and transpeptidase contained the highest number of substitutions, with the highest count in residue T245 (n = 532). As expected, no indels or premature stop codons were observed due to the essentiality of PBP2x to the bacterial peptidoglycan metabolism.

**Figure 4.**
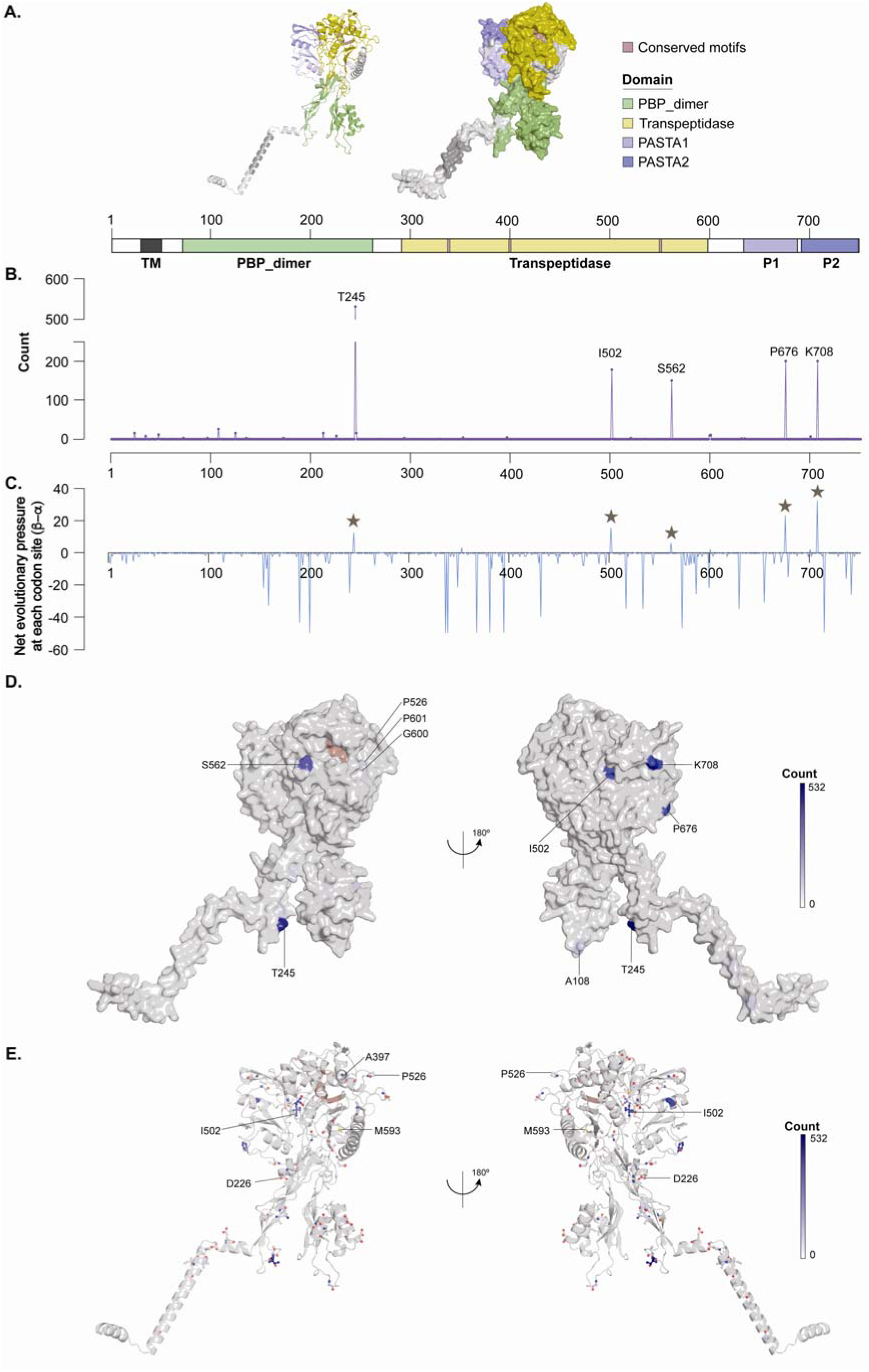
Domain architecture, substitution frequency, and site-specific selection pressure in PBP2x from *S. pyogenes*. (**A**) Ribbon and surface models of an AlphaFold-predicted structure of *S. pyogenes* PBP2x (top) and its corresponding domain schematic (bottom) coloured by domains: PBP_dimer (green), transpeptidase (yellow), PASTA 1 (P1, light purple), and PASTA 2 (P2, dark purple). The N-terminal transmembrane region (TM) is shown in dark grey. Conserved catalytic motifs within the transpeptidase domain are indicated by pink shading. (**B**) Frequency of amino acid substitutions across the PBP2x sequence among the 2,970 *S. pyogenes* genomes, aligned to the domain schematic in (A). The x-axis indicates residue position (1-751); the y-axis indicates the number of genomes carrying a substitution at each position. (**C**) Site-specific estimates of non-synonymous (β) minus synonymous (α) substitution rates calculated using FUBAR, plotted across the same sequence positions as in (B). Positive β–α values indicate codon sites with evidence of diversifying (positive) selection; negative β–α values indicate sites under purifying selection. The brown stars mark the five substitution hotspots identified in (B) (T245, I502, S562, P676, K708), all of which correspond to positive β–α values with statistical significance (P(α < β) > 0.9). (**D**) Substitution frequencies mapped onto the surface of a predicted structure of *S. pyogenes* PBP2x, shown in two orientations related by a 180° rotation about the vertical axis. Surface colouring reflects the substitution count, from white (n = 0) to dark blue (n = 532), per the scale bar. Labelled residues (T245, I502, S562, G600, P601, P676, K708) correspond to positions of high substitution frequency. The pink active-site motif is shown for reference. (**E**) Substitution frequencies mapped onto the cartoon of a predicted structure of *S. pyogenes* PBP2x, shown in two orientations related by a 180° rotation about the vertical axis. Cartoon colouring is similar to panel (D). Residues with at least one substitution are represented by balls and sticks.

*S. pyogenes emm*1 has been a dominant circulating lineage linked to global increases in invasive *S. pyogenes* infections and scarlet fever surges, including in the UK^31^. Similar to its low AMR gene content, we found only two genomes with a R632H substitution in PBP2x (Supplementary Table 1). As R632 is located in an interdomain linker, substitution at this residue is unlikely to alter MIC values for β-lactam antibiotics.

Upon mapping PBP2x substitutions onto the phylogenetic tree of our dataset, we noted seven *emm-* specific PBP2x substitution haplotypes (Figure 2, Table 1). The highest number of recurring PBP2x substitution was T245A, which could be found in all our *emm*3 genomes (Figure 2, Supplementary Table 1). Moreover, PBP2x T245A is also present in the common *emm*3 reference genome, MGAS315. To confirm the association between PBP2x T245A and *emm*3 isolates, we examined two additional datasets comprising of both invasive and non-invasive isolates collected in the UK^40^ and Australia^41^ during a similar timeframe. Within these 349 *emm*3 genomes, 99.7% (n = 348) harboured the T245A substitution in PBP2x (Supplementary Table 2, Supplementary Table 3).

**Table 1.** Seven types of *emm-*specific PBP2x substitutions. Only *emm* types with at least 5 isolates were considered.

| *emm* | PBP2x substitutions | Count |
| --- | --- | --- |
| 3 | T245A | 526/526 |
| 89 | S562T | 132/132 |
| 6 | I502V, P676S, K708E | 117/117 |
| 73 | S562T, P676S, K708E | 17/17 |
| 68 | D226N | 9/9 |
| 78 | P676S, K708E | 7/7 |
| 101 | A397V, I502V | 5/5 |

Additionally, all *emm*89 genomes contained a S562T substitution in PBP2x (Figure 2, Supplementary Table 1). Similar to the case of *emm*3/PBP2x T245A, two *emm*89 reference genomes, H293 and MGAS27061, also carried this S562T substitution in PBP2x. Additional datasets focusing on *emm*89 genomes (from both invasive and non-invasive isolates in the UK^42^ and Australia^41^) also confirmed the presence of S562T in PBP2x (Supplementary Table 2, Supplementary Table 4). However, such an association between specific PBP2x substitutions and *emm* types requires cautious interpretation as the expansion dynamics of *emm* lineages differ across countries and continents.

To estimate the selective pressure at each PBP2x codon site, we examined the mean posterior non-synonymous mutation rate (β) and the mean posterior synonymous mutation rate (α) of each of the 751 codon sites. Coupled with posterior probability of 0.9, the net evolutionary pressure at each codon site (β – α) denotes the direction of the pressure (positive vs negative)^43^. Within this dataset, five PBP2x codon sites (T245, I502, S562, P676, and K708) were under positive selection [β – α > 0, P(α < β) > 0.9; Figure 4C], indicating that non-synonymous mutations have accumulated more rapidly than neutral background drift. The association of these five positively selected codons with specific *emm* types (Table 2) implies that these corresponding substitutions in PBP2x were largely driven by *emm* lineage expansion in England and Wales prior to 2016^40,44^ rather than driven by a change in β-lactam antibiotic binding. In contrast, 31 PBP2x codons were under purifying selection [β – α < 0, P(α > β) > 0.9], and a summary of the estimated evolutionary pressure for each codon is described in Supplementary Table 5.

**Table 2.**
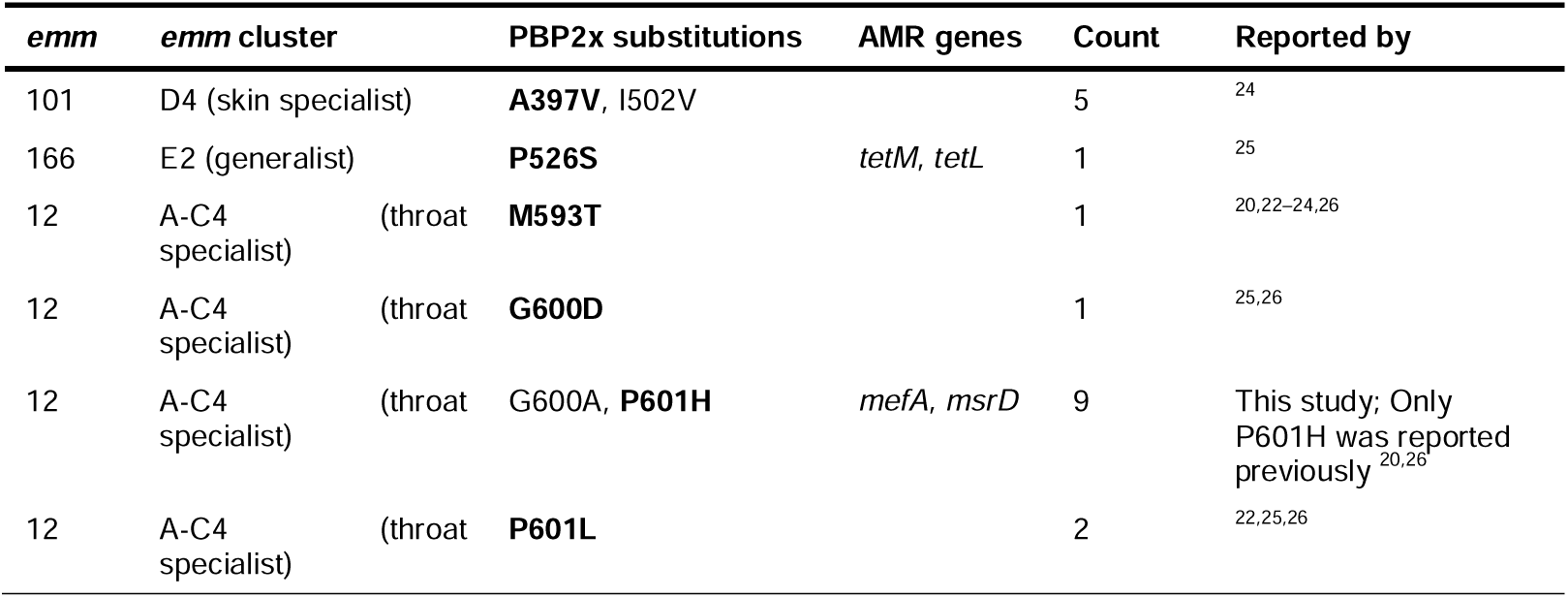
PBP2x substitutions in our dataset that have been previously associated with elevated MIC values for β-lactam antibiotics.

None of these *emm*-specific PBP2x substitutions (Table 1), except A397V, structurally maps close to the active site of PBP2x, where β-lactam antibiotics bind (Figure 4D, Figure 4E). Therefore, it is unlikely that these substitutions are associated with increased MIC values for β-lactam antibiotics. Previous studies also confirmed that *S. pyogenes* isolates with T245A (of *emm*81)^24^ or T245I (of *emm*28)^25^ did not show any MIC increases. Similarly, isolates with PBP2x S562T did not show changes in MICs^20,24^. Multiple substitutions such as I502V, P676S, K708E in PBP2x also did not change the MICs for an *emm*81 isolate^24^. In the future, continuous monitoring of PBP2x variants across *emm* types will reveal patterns of mutation accumulation and spread.

For PBP2x A397V identified in *emm*101 genomes, its proximity to the conserved motif S_399_SN (and hence, the active site) explains its association with increased MIC values for β-lactam antibiotics^24^. Within this dataset, almost all PBP2x substitutions that have been previously shown to be associated with elevated MIC values are summarised in Table 2. These PBP2x mutants were scattered across the phylogenetic tree rather than clustering in a single lineage (Figure 2), suggesting independent acquisition events in the past. In addition, we discovered nine genomes carrying a novel double PBP2x substitutions, namely G600A_P601H (Table 2). Notably, all nine genomes with PBP2x G600A_P601H also harboured the *mefA* and *msrD* genes, which confer macrolide resistance to the bacterium. No other PBP2x substitutions associated with reduced susceptibility to β-lactam antibiotics (Table 2) were found within the 145 genomes with macrolide resistance genes (Supplementary Table 1, Figure 3).

Similar to G600D^25,26^ and P601L^20,26^, we hypothesised that the double substitutions G600A_P601H in PBP2x would confer reduced susceptibility to various β-lactam antibiotics (i.e., elevated MICs) for the bacterial isolates. Broth microdilution assays and antibiotic gradient strips were used to obtain the MIC values for eight invasive isolates carrying G600A and P601H substitutions in PBP2x (Table 3). All eight isolates of PBP2x G600A_P601H showed elevated MICs for ampicillin, amoxicillin, and cefuroxime in comparison to an *emm*12 isolate without any PBP2x substitution. In particular, the MICs for penicillin were less than 1 doubling dilution away from the resistance breakpoint (≥0.03 µg/mL) defined by EUCAST^28^. Erythromycin resistance in these isolates was also verified experimentally (Supplementary Table 6). Furthermore, we tested the antibiotic susceptibility of isolates *emm*12/PBP2x M593T, *emm*12/PBP2x P601L, and *emm*1/PBP2x G600D. Consistent with others’ observations, these isolates also showed elevated MIC values for various β-lactam antibiotics, regardless of the different *emm* types across multiple studies^20,22–26^.

**Table 3.** MIC values for various β-lactam antibiotics of select PBP2x mutants. Each row represents one *S. pyogenes* isolate. Elevated MIC values are in bold.

| emm | PBP2x variant | Broth microdilution assays |  |  |  |  |  | Gradient strip tests |  |  |  |  |  |  |  |
| --- | --- | --- | --- | --- | --- | --- | --- | --- | --- | --- | --- | --- | --- | --- | --- |
| | | Ampicillin ($\mu\text{g/mL}$) | | Amoxicillin ($\mu\text{g/mL}$) | | Cefuroxime ($\mu\text{g/mL}$) | | Penicillin ($\mu\text{g/mL}$) | | Cefoxitin ($\mu\text{g/mL}$) | | Cefotaxime ($\mu\text{g/mL}$) | | Meropenem ($\mu\text{g/mL}$) | |
|  |  | Modal MIC | MIC range | Modal MIC | MIC range | Modal MIC | MIC range | Modal MIC | MIC range | Modal MIC | MIC range | Modal MIC | MIC range | Modal MIC | MIC range |
| 12 | Reference | 0.024 | 0.016-0.024 | 0.024 | 0.016-0.024 | 0.016 | 0.016-0.024 | 0.006-0.008 | 0.006-0.008 | 0.75 | 0.75 | 0.012-0.016 | 0.012-0.016 | 0.003-0.004 | 0.003-0.004 |
| 12 | M593T | 0.024 | 0.024 | **0.032** | 0.032-0.048 | 0.016 | 0.016-0.024 | **0.016** | 0.016 | 0.75-1.0 | 0.75-1.0 | 0.016-0.023 | 0.016-0.023 | **0.006** | 0.006 |
| 12 | P601L | **0.032** | 0.024-0.032 | **0.048** | 0.032-0.048 | **0.032** | 0.032 | **0.023** | 0.023 | **1.0** | 1.0 | **0.023-0.032** | 0.023-0.032 | **0.012** | 0.012 |
| 12 | G600A, P601H | **0.048** | 0.048 | **0.048** | 0.048 | **0.032** | 0.032 | **0.016-0.023** | 0.016-0.023 | 0.38-0.50 | 0.38-0.50 | 0.016 | 0.016 | **0.006** | 0.006 |
| 12 | G600A, P601H | **0.048** | 0.048 | **0.048** | 0.048 | **0.032** | 0.032 | **0.016** | 0.016 | 0.38-0.50 | 0.38-0.50 | 0.016 | 0.016 | 0.004-0.006 | 0.004-0.006 |
| 12 | G600A, P601H | **0.032** | 0.032-0.048 | **0.048** | 0.048 | 0.024 | 0.024 | not tested |  | not tested |  | not tested |  | not tested |  |
| 12 | G600A, P601H | **0.048** | 0.032-0.048 | **0.048** | 0.048 | **0.032** | 0.032 | not tested |  | not tested |  | not tested |  | not tested |  |
| 12 | G600A, P601H | **0.032** | 0.032-0.048 | **0.048** | 0.048 | **0.032** | 0.024-0.032 | not tested |  | not tested |  | not tested |  | not tested |  |
| 12 | G600A, P601H | **0.048** | 0.048 | **0.048** | 0.032-0.048 | **0.032** | 0.032 | not tested |  | not tested |  | not tested |  | not tested |  |
| 12 | G600A, P601H | **0.048** | 0.048 | **0.048** | 0.048 | **0.032** | 0.032 | not tested |  | not tested |  | not tested |  | not tested |  |
| 12 | G600A, P601H | **0.048** | 0.048 | **0.048** | 0.048 | **0.032** | 0.032 | not tested |  | not tested |  | not tested |  | not tested |  |
| 12 | I42F | 0.024 | 0.024 | 0.024 | 0.024 | 0.016 | 0.016-0.024 | 0.008-0.012 | 0.008-0.012 | 0.5-0.75 | 0.5-0.75 | 0.008-0.012 | 0.008-0.012 | **0.006** | 0.006 |
| 12 | A108T | 0.016 | 0.016-0.024 | 0.024 | 0.024-0.032 | 0.016 | 0.016 | 0.008-0.012 | 0.008-0.012 | 0.75 | 0.75 | 0.008 | 0.008 | 0.004 | 0.004 |
| 1 | Reference | 0.024 | 0.016-0.024 | 0.024 | 0.024 | 0.016 | 0.016 | 0.008 | 0.006-0.012 | 0.5 | 0.5 | 0.012-0.016 | 0.012-0.016 | 0.002-0.003 | 0.002-0.003 |
| 1 | G600D | 0.024 | 0.024 | **0.032** | 0.032 | **0.024** | 0.024 | 0.012-0.016 | 0.012-0.016 | 0.5-0.75 | 0.5-0.75 | **0.023** | 0.016-0.023 | **0.004-0.006** | 0.004-0.006 |

For broth microdilution assays, at least three independent replicates were tested for each isolate.
For gradient strip tests, two independent replicates were tested for each isolate.

The major grouping of *emm* clusters (A-C, D, and E) are associated with tissue tropisms^45^. Among our 996 genomes with PBP2x substitutions, 689 and 289 genomes belonged to the *emm* cluster A-C (throat specialists) and *emm* cluster E (generalists), respectively. *Emm* cluster D (skin specialists) was under-represented (n = 18) in general, potentially highlighting the host niches favourable for the propagation of PBP2x substitutions.

### Fluoroquinolones

Although fluoroquinolones do not constitute the standard care for infections caused by *S. pyogenes,* non-synonymous mutations in the quinolone-resistance-determining region (QRDR) of the bacterial DNA gyrase (GyrA) and topoisomerase IV (ParC) can result in fluoroquinolone non-susceptibility in this bacterium. Among the 2,970 genomes analysed in this study, we did not detect any non-synonymous mutations from residues 81 to 85 in the QRDR of GyrA. However, 151 genomes carried non-synonymous mutations from residues 79 to 83 in the QRDR of ParC, including S79F, S79A, S79Y, and D83G, which previously resulted in fluoroquinolone resistance in *S. pyogenes*^46–48^. None of these 151 genomes encoded PBP2x substitutions associated with reduced susceptibility to β-lactam antibiotics (Table 2).

## DISCUSSION

Our study provides a comprehensive investigation of the prevalence and diversity of AMR determinants in 2,970 invasive *S. pyogenes* isolates from England and Wales (2001 to 2015), with a particular focus on bacterial susceptibility to β-lactam antibiotics. Although no true clinical resistance to β-lactam antibiotics has been observed, penicillin-resistant *S. pyogenes* expressing penicillin-binding proteins (PBPs) with low affinity for β-lactam antibiotics have previously been isolated in the laboratory under the right selective pressures^49,50^, demonstrating the biological feasibility of β-lactam resistance in *S. pyogenes*. The emergence of *S. pyogenes* isolates showing elevated MICs for β-lactam antibiotics in other countries has been concerning^19,20,22–26^, and whether this threat existed in the UK was unclear. Overall, this study has resulted in three key insights:

### (i) AMR genes clustered in non-dominating emm types

Within *S. pyogenes*, many AMR genes (such as *tetM*, *ermA*, and *ermB*) spread within clonal lineages via mobile genetic elements (prophages, integrative conjugative elements, transposons) rather than being distributed evenly across the various *emm* types. Despite the small proportion of genomes with acquired AMR genes (10.6%) in this study, the concentration of AMR genes in non-dominating *emm* types at that time aligns with this expectation. It is unclear whether dominating lineages such as *emm*1 and *emm*12 remain unchallenged by these mobile genetic elements or whether these dominating lineages have specific genetic barriers that prevented incorporation of AMR genes (e.g., lower competence in *emm*1 in comparison to *emm*49^51^). Epidemiological studies indicated that the populations where non-dominating *emm* types have been more prevalent tend to be marginalised groups, possibly indicating a prior treatment-based selection pressure, where oral regimens, such as doxycycline, can be preferred treatment practices in these populations^52^. Distinguishing these two scenarios will require longitudinal, unbiased sampling of *S. pyogenes* to understand the interplay between resistance carriage and lineage dominance.

### (ii) Some sequence variations in PBP2x appeared to be emm type-specific

We found seven *emm*-specific combinations of PBP2x substitutions that appeared to have been fixed within lineages. Five *emm*-associated substitutions, namely T245A, S562T, I502V, P676S, and K708E, were under positive selection but without any association with elevated MICs for β-lactam antibiotics, implying that these substitutions – at least within this study – were driven by *emm* lineage expansion during the sampling period. A similar observation was also reported previously in which 98% of *emm*12 isolates from Iceland were found to harbour the M593T substitution in PBP2x^23^. Further evidence is required to understand if these substitutions provide any selective advantage to the lineages in England and Wales. Moreover, the existing PBP2x diversity in our dataset raises the question whether these sequences will evolve in different trajectories (i.e., accumulating different sets of mutations in the future), which subsequently might facilitate the emergence of β-lactam non-susceptibility in *S. pyogenes*.

### (iii) Co-occurrence of a novel PBP2x G600A_P601H mutant with macrolide resistance genes in nine emm12 isolates

Among the 145 macrolide-resistant isolates, 9 *emm*12 isolates were of particular concern as they showed elevated MICs for various β-lactam antibiotics. Most of these isolates were identified in the same geographical area, suggestive of localised transmission. Within these isolates, the co-occurrence of G600A and P601H in PBP2x was novel, although both substitutions have been reported individually and P601H has been associated with elevated MICs for β-lactam antibiotics^20,25,26^. Both G600 and P601 lie at the beginning of helix α11, which is approximately 10 Å from the catalytic motifs, indicating an unlikelihood of these residues directly participating in substrate or ligand binding. Yet, G600 and P601 sit above the β3 strand on the lower lid of the substrate cavity, suggesting that they might influence β3 or the conformation of the cavity lid. In five (out of six) cefotaxime-resistant isolates of *S. pneumoniae*, G597 (the equivalent of P601) was replaced by an aspartic acid and this substitution contributed to the second step of the development of cefotaxime resistance^53^. Another cefotaxime-resistant isolate of *S. pneumoniae* variant carrying substitutions in three residues (T550, S596, and G601, which correspond to T553, G600, and Q605 in *S. pyogenes*) showed 30-fold higher MIC for cefotaxime and 300-fold lower acylation efficiency when compared to a wildtype isolate without the substitutions^17^. Coupled with previous increases in MICs for β-lactam antibiotics shown by G600 and P601 mutants of *S. pyogenes*^22,25,26^, it is likely that G600 and P601 are important determinants of the bacterial susceptibility to β-lactam antibiotics.

Further investigation is required to understand the fitness of these PBP2x mutants and their infection outcomes. Although Gutmann & Tomasz (1982) reported that *pbp* mutations in penicillin-resistant *S. pyogenes* (induced in the laboratory) could incur a high fitness cost^50^, recent evidence is suggesting a more nuanced picture. Notably, a PBP2x P601L mutant of *S. pyogenes* displayed a higher *in vivo* fitness than an isogenic wildtype during subtherapeutic penicillin therapy in mice with necrotizing myositis^21^. The PBP2x T553K mutants also did not show growth differences^19^, suggesting that not all *pbp2x* mutations are deleterious to the bacterium. Furthermore, mutations elsewhere in the bacterial genomes could compensate for deleterious *pbp* mutations, but their identity and their compensatory mechanisms remain unclear.

The development of β-lactam non-susceptibility in *S. pneumoniae* is a multifactorial process, frequently involving genetic alterations in *pbp* and non-*pbp* genes^38^. In addition, other key streptococcal pathogens that have evolved a significant degree of non-susceptibility to β-lactam antibiotics include *Streptococcus agalactiae*^54^, *Streptococcus dysgalactiae* subsp. *equisimilis*^55^, and *Streptococcus suis*^56^ – all of which carried *pbp* mutations with low affinity for β-lactam antibiotics. Do we expect *S. pyogenes* to evolve non-susceptibility to β-lactam antibiotics in the same way as these streptococcal pathogens did? The answer is unclear for *S. pyogenes,* as we have yet to observe high level non-susceptibility in *S. pyogenes*. However, we and others have captured the small MIC changes in *S. pyogenes* with select PBP2x substitutions, which could be the first step in the development of β-lactam non-susceptibility. Discerning which *pbp2x* mutations are important for this development, and which ones are not, is critical for future predictions.

In our attempt to understand the AMR landscape in *S. pyogenes*, this study is limited by the following factors: (i) A gap in the collection years between 2012 and 2013; (ii) a lack of regional variation (our dataset was derived only from England and Wales); (iii) a lack of non-invasive *S. pyogenes* isolates from the same geographic locations for comparative purposes; and (iv) a potential lack of genotypic diversity as only 74 *emm* types (of all 275 *emm* types) were covered. Moving forward, we expect that a larger and more recent bacterial genomics dataset coupled with clinical and epidemiological data would enable a more contextualised understanding of the evolving AMR landscape in *S. pyogenes*.

Rare *emm* types such as *emm*83 have been disproportionately affecting inclusion health populations, particularly people experiencing homelessness and people who inject drugs^32,57^. Our study showed that these less frequently encountered *emm* types carried a greater burden of acquired AMR genes. Together, these findings highlight the importance of agile adjustment of antimicrobial therapy that is evidence-based, incorporating local molecular epidemiology, temporal trends in *emm* type distribution, AMR patterns, and patient-level risk factors into treatment decisions. Such considerations may be particularly relevant for patients with penicillin allergy, for whom reliance on alternative agents may be affected by the higher prevalence of resistance observed in certain *emm* lineages, although confirmation of reported penicillin allergy should also be considered whenever feasible.

*S. pyogenes* infections show an unequal disease burden with a disproportionate impact on low socioeconomic communities including the elderly and inclusion health populations^10,32^. Within England and Wales, a higher transmission rate of *S. pyogenes* is possible with the changing demographics and socioeconomics in the community, which in turn may encourage the spread of AMR determinants, including *pbp2x* mutations. Ongoing lineage diversification of *S. pyogenes* and the co-occurrence of AMR determinants therefore warrant continuous genomic surveillance and further antibiotic susceptibility testing for *S. pyogenes*.

## METHODS

### Bacterial genomic datasets

A total of 4,256 Illumina whole-genome sequencing reads of *S. pyogenes* isolates^29,30,40–42^ were downloaded from the European Nucleotide Archive (www.ebi.ac.uk/ena/browser/home). Table 4 summarises the information of these genomes.

**Table 4.** List of invasive *S. pyogenes* genomes analysed in this study.

| Dataset | Number of genomes analysed in this study* | Collection years (duration) | Locations | Invasiveness | ENA Project Accession |
| --- | --- | --- | --- | --- | --- |
| <u>Datasets used for this study</u> |  |  |  |  |  |
| British Society for Antimicrobial Chemotherapy (BSAC) Bacteraemia Resistance Surveillance Program<sup>29</sup> | 318 (344) | 2001 to 2011 (11 years) | England and Wales | Invasive | PRJEB4679 |
| Public Health England National Streptococcal Reference Laboratory<sup>30</sup> | 2,652 (3,047) | April 2014 to May 2015 (14 months) | England and Wales | Invasive | PRJEB17673 |
| <u>Comparator datasets</u> |  |  |  |  |  |
| *emm*3<sup>40</sup> | 332 (450) | 2001 to 2013 (13 years) | England | Invasive + Non-invasive | PRJEB2437 |
| *emm*89<sup>42</sup> | 85 (97) | 2004 to 2013 (10 years) | UK | Invasive | PRJEB4679 |
| Australian isolates<sup>41</sup> | 281 (318) | 2007 to 2017 (11 years) and 2019-2021 (3 years) | Australia | Invasive + Non-invasive | PRJNA996294 |

\*The number of genomes reported by the original study is in brackets.

### Quality control and assembly of bacterial genomes

Adaptor sequences and low-quality bases were removed from the raw sequencing reads using Trimmomatic (v0.39)^58^ prior to species confirmation using Kraken2 (v2.1.43)^59^ and Bracken (v2.9)^60^. Each bacterial genome was assembled *de novo* using SPAdes (v3.15.5)^61^, with the quality of each contig assessed using Quast (v5.2.0)^62^ and CheckM2 (v1.2.0)^63^. Contigs of <200 bp were excluded from downstream analysis. Assembled genomes were annotated using Bakta (v1.9.3)^64^ with its full database v5.1.

### Genotyping and screening of antimicrobial resistance determinants

Multilocus sequence typing (MLST) was performed on draft genome assemblies using *mlst* (v2.23.0; github.com/tseemann/mlst) with the reference database of *S. pyogenes* on PubMLST (pubmlst.org/organisms/streptococcus-pyogenes). The *emm* type and tissue tropism types were assigned with *emmtyper* (v0.2.0; github.com/MDU-PHL/emmtyper). Antimicrobial resistance genes in each genome were identified using ABRicate (v1.0.1; github.com/tseemann/abricate) based on the ResFinder database (last updated 04/11/2023)^65^. Mutations within the *pbp*, *gyrA*, and *parC* genes were identified using megaBLAST implemented in *rasti* (v0.0.1; github.com/wanyuac/rasti), and were compared against the corresponding alleles from an *emm*1 reference strain *S. pyogenes* MGAS5005 (Genbank accession CP000017.2). Other reference genomes used in this study: *emm*3 MGAS315 (Genbank accession: AE014074.1); *emm*4 MGAS10750 (Genbank accession: CP000262); *emm*6 MGAS10394 (Genbank CP000003.1); *emm*11 (Genbank accession: CP035432.1); *emm*12 HKU12 (Genbank accession: AFRY01000001.1); *emm*28 MGAS6180 (Genbank accession: CP000056.2); *emm*75 (Genbank accession: CP033621.1); *emm*89 H293 (Genbank accession: HG316453.2) and *emm*89 MGAS27061 (Genbank accession: CP013840.1).

### Phylogenetic analysis

Single nucleotide polymorphisms (SNPs) from the core genome set were identified using Snippy (v4.6.0; github.com/tseemann/snippy), and any recombinant regions were removed using Gubbins (v3.3.1)^66^. A pangenome was constructed from the annotated draft genome assemblies using Panaroo (v1.51.1)^67^. A species-wide tree was reconstructed using IQ-TREE with the GTR+G model and 1,000 bootstrap replicates (v2.3.3)^68^ and was visualised using iTOL v7.0^69^. SNPs between genome pairs were assessed using snp-dists (v0.8.2; github.com/tseemann/snp-dists).

### Evaluation of PBP2x variants

Protein structures were visualised using PyMOL^70^, and PBP2x substitutions were rendered onto a predicted structure of *S. pyogenes* PBP2x (AlphaFold Protein Structure Database^71^ accession: AF-Q99YK1-F1). The FUBAR method from the HyPhy package (v1.3.1)^71^ was used to assess the strength of selection pressure on each codon site.

### Antibiotic susceptibility assays

Bacterial susceptibility to penicillin, cefotaxime, cefoxitin, and meropenem was determined using gradient strip testing (Liofilchem MIC Test Strips) according to the manufacturer’s instructions. Briefly, individual colonies of *S. pyogenes* grown on Columbia Blood Agar (E&O Laboratories) were suspended in 0.85% saline and their turbidity was adjusted to 0.5 McFarland standard (equivalent to 10^6^ cfu/mL). Using a sterile swab, each bacterial suspension was spread onto a Mueller-Hinton agar with 5% horse blood and 20 mg/L nicotinamide adenine dinucleotide (E&O Laboratories). The relevant MIC test strip was placed on top of the bacterial lawn, and the agar plates were incubated at 37°C in the presence of 5% CO_2_ for 18 hours. The minimum inhibitory concentration (MIC) was read from the scale at the intersection where the growth inhibition intersects the MIC test strip.

MICs for amoxicillin, ampicillin, and cefuroxime were determined using the broth microdilution assays^72^. A combination of two-fold serial dilutions and half-step dilutions of an antibiotic was set up in 96-well microtiter plate, with each well inoculated with 10^5^ bacterial cells. All wells contained a final volume of 150 µL. For each antibiotic, the MIC was defined as the lowest concentration of an antibiotic that prevented bacterial growth (i.e., OD_600_ <0.1) after 18 hours of incubation at 37°C and 5% CO_2_.

All MIC results were interpreted according to the CLSI (34^th^ ed.)^28^ and the EUCAST v16.0^27^ guidelines. CLSI specifies the resistance breakpoints as follows: penicillin ≥0.12 µg/mL; ampicillin ≥0.25 µg/mL; cefotaxime ≥0.5 µg/mL; meropenem ≥0.5 µg/mL; not defined for amoxicillin, cefuroxime, and cefoxitin^28^. EUCAST defines the resistance breakpoint for penicillin as ≥0.03 µg/mL; whereas the susceptibility of *S. pyogenes* to other penicillins (ampicillin, amoxicillin), cephalosporins (cefotaxime, cefuroxime, cefoxitin) and carbapenems (meropenem) is inferred from the susceptibility of benzylpenicillin^27^. The *pbp2x* mutations of all tested *S. pyogenes* isolates were verified by Sanger sequencing (Azenta).

Bacterial susceptibility to erythromycin and clindamycin was tested using disk diffusion assays. In each test, an erythromycin disk (15 µg) and a clindamycin disk (2 µg) were placed 12 mm apart (edge to edge) on a bacterial lawn produced as described for the gradient strip testing. The diameter of inhibition zone was read using callipers after an incubation at 37°C and 5% CO_2_ for 18 hours. CLSI defines the resistance breakpoints for erythromycin as ≤15 mm and clindamycin as ≤15 mm^28^. EUCAST defines the resistance breakpoints for erythromycin as <21 mm and clindamycin as <17 mm^28^.

## AUTHOR CONTRIBUTIONS

GS: data curation, formal analysis, investigation, visualization, and writing – original draft. ALM: formal analysis and investigation. SG: data curation and investigation. RLG: formal analysis. YW: software. AV: investigation. JC and KM: resources. SS: resources. EJ: conceptualization, data curation, formal analysis, funding acquisition, methodology, supervision, project administration, and writing – original draft. VWCS: conceptualization, data curation, formal analysis, funding acquisition, methodology, supervision, project administration, visualization, and writing – original draft. All authors reviewed and edited the writing.

## ACKNOWLEDGEMENTS

This work was supported by an MRC Career Development Award (MR/X007421/1) to VWCS. EJ acknowledges funding from Genesis Research Trust. YW is funded by the David Price Evans Endowment (UGG10057) at the University of Liverpool. The authors acknowledge the support of the NIHR Biomedical Research Centre awarded to Imperial College London. GS, YW, SS, and EJ were affiliated with the NIHR Health Protection Research Unit in Healthcare Associated Infections and Antimicrobial Resistance at Imperial College London in partnership with the UKHEA, in collaboration with Imperial Healthcare Partners, University of Cambridge, and University of Warwick (NIHR200876). The views expressed in this article are those of the authors and not necessarily of the NHS, the NIHR, or the Department of Health and Social Care.

## DATA AVAILABILITY

This study only used publicly available Illumina whole-genome sequencing data of *S. pyogenes* downloaded from the European Nucleotide Archive (www.ebi.ac.uk/ena/browser/home). The authors confirm all supporting data including the run accessions of sequencing reads have been provided within the article or through supplementary files.

