## Supplementary Data for "Diversity of penicillin-binding protein 2x in *Streptococcus pyogenes* from England and Wales, 2001 to 2015"

**Supplementary Data. All 43 PBP2x sequence variants (including the reference PBP2x sequence) detected in this study.**

>PBP2x

MKKWQKYVLDYVVRDRRTPVENRVRVGQNMMLLTIFIFFIFIINFMIIIGTDQKFGVSLSEGAKKVYQETVTIQAKRGTIYDRNGTAIAVDSTTYSIYAILDKSFVSASDEKLYVQPSQYETVADILKKHLGMKKTDVIKQLKRKGLFQVSFGPSGSGISYSTMSTIQKAMEDAKIKGIAFTTSPGRMYPNGTFASEFIGLASLTEDKKTGVKSLVGKTGLEASFDKILSGQDGVITYQKDRNGTTLLGTGKTVKKAIDGKDIYTTLSEPIQTFLETQMDVFQAKSNGQLASATLVNAKTGEILATTQRPTYNADTLKGLENTNYKWYSALHQGNFEPGSTMKVMTLAAAIDDKVFNPNETFSNANGLTIADATIQDWSINEGISTGQYMNYAQGFAFSSNVGMTKLEQKMGNAKWMNYLTKFRFGFPTRFGLKDEDAGIFPSDNIVTQAMSAFGQGISVTQIQMLRAFTAISNNGEMLEPQFISQIYDPNTASFRTANKEIVGKPVSKKAASETRQYMIGVGTDPEFGTLYSKTFGPIIKVGDLPVAVKSGTAQIGSEDGSGYQDGGLTNYVYSVVAMVPADKPDFLMYVTMTKPQHFGPLFWQDVVNPVLEEAYLMQDTLTKPVVSDANRQTTYKLPNFVGKNPGETSSELRRNLVQPVVLGTGSKIKKVSHQPGQTLTENQQVLILSDRFVEVPDMYGWTKSNVKTFAKWTGIDISFKGTDSGRVMKQSVDVGKSLKKIKKMTITLGD*

>PBP2x.41

MKKWQKYVLDYVVRDRRTPVENRVRVGQNMMLLTTFIFFIFIINFMIIIGTDQKFGVSLSEGAKKVYQETVTIQAKRGTIYDRNGTAIAVDSTTYSIYAILDKSFVSASDEKLYVQPSQYETVADILKKHLGMKKTDVIKQLKRKGLFQVSFGPSGSGISYSTMSTIQKAMEDAKIKGIAFTTSPGRMYPNGTFASEFIGLASLTEDKKTGVKSLVGKTGLEASFDKILSGQDGVITYQKDRNGTTLLGTGKTVKKAIDGKDIYTTLSEPIQTFLETQMDVFQAKSNGQLASATLVNAKTGEILATTQRPTYNADTLKGLENTNYKWYSALHQGNFEPGSTMKVMTLAAAIDDKVFNPNETFSNANGLTIADATIQDWSINEGISTGQYMNYAQGFAFSSNVGMTKLEQKMGNAKWMNYLTKFRFGFPTRFGLKDEDAGIFPSDNIVTQAMSAFGQGISVTQIQMLRAFTAISNNGEMLEPQFISQIYDPNTASFRTANKEVVGKPVSKKAASETRQYMIGVGTDPEFGTLYSKTFGPIIKVGDLPVAVKSGTAQIGSEDGSGYQDGGLTNYVYSVVAMVPADKPDFLMYVTMTKPQHFGPLFWQDVVNPVLEEAYLMQDTLTKPVVSDANRQTTYKLPNFVGKNPGETSSELRRNLVQPVVLGTGSKIKKVSHQSGQTLTENQQVLILSDRFVEVPDMYGWTKSNVETFAKWTGIDISFKGTDSGRVMKQSVDVGKSLKKIKKMTITLGD*

>PBP2x.25

MKKWQKYVLDYVVRDRRTPVENRIRVGQNMMLLTIFIFFIFIINFMIIIGTDQKFGVSLSEGAKKVYQETVTIQAKRGTIYDRNGTAIAVDSTTYSIYAILDKSFVSASDEKLYVQPSQYETVADILKKHLGMKKTDVIKQLKRKGLFQVSFGPSGSGISYSTMSTIQKAMEDAKIKGIAFTTSPGRMYPNGTFASEFIGLASLTEDKKTGVNSLVGKTGLEASFDKILSGQDGVITYQKDRNGTALLGTGKTVKKAIDGKDIYTTLSEPIQTFLETQMDVFQAKSNGQLASATLVNAKTGEILATTQRPTYNADTLKGLENTNYKWYSALHQGNFEPGSTMKVMTLAAAIDDKVFNPNETFSNANGLTIADATIQDWSINEGISTGQYMNYAQGFAFSSNVGMTKLEQKMGNAKWMNYLTKFRFGFPTRFGLKDEDAGIFPSDNIVTQAMSAFGQGISVTQIQMLRAFTAISNNGEMLEPQFISQIYDPNTASFRTANKEIVGKPVSKKAASETRQYMIGVGTDPEFGTLYSKTFGPIIKVGDLPVAVKSGTAQIGSEDGSGYQDGGLTNYVYSVVAMVPADKPDFLMYVTMTKPQHFGPLFWQDVVNPVLEEAYLMQDTLTKPVVSDANRQTTYKLPNFVGKNPGETSSELRRNLVQPVVLGTGSKIKKVSHQPGQTLTENQQVLILSDRFVEVPDMYGWTKSNVKTFAKWTGIDISFKGTDSGRVMKQSVDVGKSLKKIKKMTITLGD*

>PBP2x.88

MKKWQKYVLDYVVRDRRTPVENRVRVGQNMMLLTIFIFFIFFINFMIIIGTDQKFGVSLSEGAKKVYQETVTIQAKRGTIYDRNGTAIAVDSTTYSIYAILDKSFVSASDEKLYVQPSQYETVADILKKHLGMKKTDVIKQLKRKGLFQVSFGPSGSGISYSTMSTIQKAMEDAKIKGIAFTTSPGRMYPNGTFASEFIGLASLTEDKKTGVKSLVGKTGLEASFDKILSGQDGVITYQKDRNGTTLLGTGKTVKKAIDGKDIYTTLSEPIQTFLETQMDVFQAKSNGQLASATLVNAKTGEILATTQRPTYNADTLKGLENTNYKWYSALHQGNFEPGSTMKVMTLAAAIDDKVFNPNETFSNANGLTIADATIQDWSINEGISTGQYMNYAQGFAFSSNVGMTKLEQKMGNAKWMNYLTKFRFGFPTRFGLKDEDAGIFPSDNIVTQAMSAFGQGISVTQIQMLRAFTAISNNGEMLEPQFISQIYDPNTASFRTANKEIVGKPVSKKAASETRQYMIGVGTDPEFGTLYSKTFGPIIKVGDLPVAVKSGTAQIGSEDGSGYQDGGLTNYVYSVVAMVPADKPDFLMYVTMTKPQHFGPLFWQDVVNPVLEEAYLMQDTLTKPVVSDANRQTTYKLPNFVGKNPGETSSELRRNLVQPVVLGTGSKIKKVSHQPGQTLTENQQVLILSDRFVEVPDMYGWTKSNVKTFAKWTGIDISFKGTDSGRVMKQSVDVGKSLKKIKKMTITLGD*

>PBP2x.67

MKKWQKYVLDYVVRDRRTPVENRVRVGQNMMLLTIFIFFIFIINFMVIIGTDQKFGVSLSEGAKKVYQETVTIQAKRGTIYDRNGTAIAVDSTTYSIYAILDKSFVSASDEKLYVQPSQYETVADILKKHLGMKKTDVIKQLKRKGLFQVSFGPSGSGISYSTMSTIQKAMEDAKIKGIAFTTSPGRMYPNGTFASEFIGLASLTEDKKTGVKSLVGKTGLEASFDKILSGQDGVITYQKDRNGATLLGTGKTVKKAIDGKDIYTTLSEPIQTFLETQMDVFQAKSNGQLASATLVNAKTGEILATTQRPTYNADTLKGLENTNYKWYSALHQGNFEPGSTMKVMTLAAAIDDKVFNPNETFSNANGLTIADATIQDWSINEGISTGQYMNYAQGFAFSSNVGMTKLEQKMGNAKWMNYLTKFRFGFPTRFGLKDEDAGIFPSDNIVTQAMSAFGQGISVTQIQMLRAFTAISNNGEMLEPQFISQIYDPNTASFRTANKEIVGKPVSKKAASETRQYMIGVGTDPEFGTLYSKTFGPIIKVGDLPVAVKSGTAQIGSEDGSGYQDGGLTNYVYSVVAMVPADKPDFLMYVTMTKPQHFGPLFWQDVVNPVLEEAYLMQDTLTKPVVSDANRQTTYKLPNFVGKNPGETSSELRRNLVQPVVLGTGSKIKKVSHQPGQTLTENQQVLILSDRFVEVPDMYGWTKSNVKTFAKWTGIDISFKGTDSGRVMKQSVDVGKSLKKIKKMTITLGD*

>PBP2x.21

MKKWQKYVLDYVVRDRRTPVENRVRVGQNMMLLTIFIFFIFIINFMIVIGTDQKFGVSLSEGAKKVYQETVTIQAKRGTIYDRNGTAIAVDSTTYSIYAILDKSFVSASDEKLYVQPSQYETVADILKKHLGMKKTDVIKQLKRKGLFQVSFGPSGSGISYSTMSTIQKAMEDAKIKGIAFTTSPGRMYPNGTFASEFIGLASLTEDKKTGVKSLVGKTGLEASFDKILSGQDGVITYQKDRNGTTLLGTGKTVKKAIDGKDIYTTLSEPIQTFLETQMDVFQAKSNGQLASATLVNAKTGEILATTQRPTYNADTLKGLENTNYKWYSALHQGNFEPGSTMKVMTLAAAIDDKVFNPNETFSNANGLTIADATIQDWSINEGISTGQYMNYAQGFAFSSNVGMTKLEQKMGNAKWMNYLTKFRFGFPTRFGLKDEDAGIFPSDNIVTQAMSAFGQGISVTQIQMLRAFTAISNNGEMLEPQFISQIYDPNTASFRTANKEIVGKPVSKKAASETRQYMIGVGTDPEFGTLYSKTFGPIIKVGDLPVAVKSGTAQIGSEDGSGYQDGGLTNYVYSVVAMVPADKPDFLMYVTMTKPQHFGPLFWQDVVNPVLEEAYLMQDTLTKPVVSDANRQTTYKLPNFVGKNPGETSSELRRNLVQPVVLGTGSKIKKVSHQPGQTLTENQQVLILSDRFVEVPDMYGWTKSNVKTFAKWTGIDISFKGTDSGRVMKQSVDVGKSLKKIKKMTITLGD*

>PBP2x.100

MKKWQKYVLDYVVRDRRTPVENRVRVGQNMMLLTIFIFFIFIINFMIIIGTGQKFGVSLSEGAKKVYQETVTIQAKRGTIYDRNGTAIAVDSTTYSIYAILDKSFVSASDEKLYVQPSQYETVADILKKHLGMKKTDVIKQLKRKGLFQVSFGPSGSGISYSTMSTIQKAMEDAKIKGIAFTTSPGRMYPNGTFASEFIGLASLTEDKKTGVKSLVGKTGLEASFDKILSGQDGVITYQKDRNGTTLLGTGKTVKKAIDGKDIYTTLSEPIQTFLETQMDVFQAKSNGQLASATLVNAKTGEILATTQRPTYNADTLKGLENTNYKWYSALHQGNFEPGSTMKVMTLAAAIDDKVFNPNETFSNANGLTIADATIQDWSINEGISTGQYMNYAQGFAFSSNVGMTKLEQKMGNAKWMNYLTKFRFGFPTRFGLKDEDAGIFPSDNIVTQAMSAFGQGISVTQIQMLRAFTAISNNGEMLEPQFISQIYDPNTASFRTANKEIVGKPVSKKAASETRQYMIGVGTDPEFGTLYSKTFGPIIKVGDLPVAVKSGTAQIGSEDGSGYQDGGLTNYVYSVVAMVPADKPDFLMYVTMTKPQHFGPLFWQDVVNPVLEEAYLMQDTLTKPVVSDANRQTTYKLPNFVGKNPGETSSELRRNLVQPVVLGTGSKIKKVSHQPGQTLTENQQVLILSDRFVEVPDMYGWTKSNVKTFAKWTGIDISFKGTDSGRVMKQSVDVGKSLKKIKKMTITLGD*

>PBP2x.97

MKKWQKYVLDYVVRDRRTPVENRVRVGQNMMLLTIFIFFIFIINFMIIIGTDQKFGVSLSGGAKKVYQETVTIQAKRGTIYDRNGTAIAVDSTTYSIYAILDKSFVSASDEKLYVQPSQYETVADILKKHLGMKKTDVIKQLKRKGLFQVSFGPSGSGISYSTMSTIQKAMEDAKIKGIAFTTSPGRMYPNGTFASEFIGLASLTEDKKTGVKSLVGKTGLEASFDKILSGQDGVITYQKDRNGTTLLGTGKTVKKAIDGKDIYTTLSEPIQTFLETQMDVFQAKSNGQLASATLVNAKTGEILATTQRPTYNADTLKGLENTNYKWYSALHQGNFEPGSTMKVMTLAAAIDDKVFNPNETFSNANGLTIADATIQDWSINEGISTGQYMNYAQGFAFSSNVGMTKLEQKMGNAKWMNYLTKFRFGFPTRFGLKDEDAGIFPSDNIVTQAMSAFGQGISVTQIQMLRAFTAISNNGEMLEPQFISQIYDPNTASFRTANKEIVGKPVSKKAASETRQYMIGVGTDPEFGTLYSKTFGPIIKVGDLPVAVKSGTAQIGSEDGSGYQDGGLTNYVYSVVAMVPADKPDFLMYVTMTKPQHFGPLFWQDVVNPVLEEAYLMQDTLTKPVVSDANRQTTYKLPNFVGKNPGETSSELRRNLVQPVVLGTGSKIKKVSHQPGQTLTENQQVLILSDRFVEVPDMYGWTKSNVKTFAKWTGIDISFKGTDSGRVMKQSVDVGKSLKKIKKMTITLGD*

>PBP2x.40

MKKWQKYVLDYVVRDRRTPVENRVRVGQNMMLLTIFIFFIFIINFMIIIGTDQKFGVSLSEGAKKVYQETVTMQAKRGTIYDRNGTAIAVDSTTYSIYAILDKSFVSASDEKLYVQPSQYETVADILKKHLGMKKTDVIKQLKRKGLFQVSFGPSGSGISYSTMSTIQKAMEDAKIKGIAFTTSPGRMYPNGTFASEFIGLASLTEDKKTGVKSLVGKTGLEASFDKILSGQDGVITYQKDRNGTTLLGTGKTVKKAIDGKDIYTTLSEPIQTFLETQMDVFQAKSNGQLASATLVNAKTGEILATTQRPTYNADTLKGLENTNYKWYSALHQGNFEPGSTMKVMTLAAAIDDKVFNPNETFSNANGLTIADATIQDWSINEGISTGQYMNYAQGFAFSSNVGMTKLEQKMGNAKWMNYLTKFRFGFPTRFGLKDEDAGIFPSDNIVTQAMSAFGQGISVTQIQMLRAFTAISNNGEMLEPQFISQIYDPNTASFRTANKEIVGKPVSKKAASETRQYMIGVGTDPEFGTLYSKTFGPIIKVGDLPVAVKSGTAQIGSEDGSGYQDGGLTNYVYSVVAMVPADKPDFLMYVTMTKPQHFGPLFWQDVVNPVLEEAYLMQDTLTKPVVSDANRQTTYKLPNFVGKNPGETSSELRRNLVQPVVLGTGSKIKKVSHQPGQTLTENQQVLILSDRFVEVPDMYGWTKSNVKTFAKWTGIDISFKGTDSGRVMKQSVDVGKSLKKIKKMTITLGD*

>PBP2x.27

MKKWQKYVLDYVVRDRRTPVENRVRVGQNMMLLTIFIFFIFIINFMIIIGTDQKFGVSLSEGAKKVYQETVTIQAKRGTIYDRNGTAIAVDSTTYSVYAILDKSFVSASDEKLYVQPSQYETVADILKKHLGMKKTDVIKQLKRKGLFQVSFGPSGSGISYSTMSTIQKAMEDAKIKGIAFTTSPGRMYPNGTFASEFIGLASLTEDKKTGVKSLVGKTGLEASFDKILSGQDGVITYQKDRNGTTLLGTGKTVKKAIDGKDIYTTLSEPIQTFLETQMDVFQAKSNGQLASATLVNAKTGEILATTQRPTYNADTLKGLENTNYKWYSALHQGNFEPGSTMKVMTLAAAIDDKVFNPNETFSNANGLTIADATIQDWSINEGISTGQYMNYAQGFAFSSNVGMTKLEQKMGNAKWMNYLTKFRFGFPTRFGLKDEDAGIFPSDNIVTQAMSAFGQGISVTQIQMLRAFTAISNNGEMLEPQFISQIYDPNTASFRTANKEIVGKPVSKKAASETRQYMIGVGTDPEFGTLYSKTFGPIIKVGDLPVAVKSGTAQIGSEDGTGYQDGGLTNYVYSVVAMVPADKPDFLMYVTMTKPQHFGPLFWQDVVNPVLEEAYLMQDTLTKPVVSDANRQTTYKLPNFVGKNPGETSSELRRNLVQPVVLGTGSKIKKVSHQPGQTLTENQQVLILSDRFVEVPDMYGWTKSNVKTFAKWTGIDISFKGTDSGRVMKQSVDVGKSLKKIKKMTITLGD*

>PBP2x.26

MKKWQKYVLDYVVRDRRTPVENRVRVGQNMMLLTIFIFFIFIINFMIIIGTDQKFGVSLSEGAKKVYQETVTIQAKRGTIYDRNGTAIAVDSTTYSIYAILDKSFVSTSDEKLYVQPSQYETVADILKKHLGMKKTDVIKQLKRKGLFQVSFGPSGSGISYSTMSTIQKAMEDAKIKGIAFTTSPGRMYPNGTFASEFIGLASLTEDKKTGVKSLVGKTGLEASFDKILSGQDGVITYQKDRNGTTLLGTGKTVKKAIDGKDIYTTLSEPIQTFLETQMDVFQAKSNGQLASATLVNAKTGEILATTQRPTYNADTLKGLENTNYKWYSALHQGNFEPGSTMKVMTLAAAIDDKVFNPNETFSNANGLTIADATIQDWSINEGISTGQYMNYAQGFAFSSNVGMTKLEQKMGNAKWMNYLTKFRFGFPTRFGLKDEDAGIFPSDNIVTQAMSAFGQGISVTQIQMLRAFTAISNNGEMLEPQFISQIYDPNTASFRTANKEIVGKPVSKKAASETRQYMIGVGTDPEFGTLYSKTFGPIIKVGDLPVAVKSGTAQIGSEDGSGYQDGGLTNYVYSVVAMVPADKPDFLMYVTMTKPQHFGPLFWQDVVNPVLEEAYLMQDTLTKPVVSDANRQTTYKLPNFVGKNPGETSSELRRNLVQPVVLGTGSKIKKVSHQPGQTLTENQQVLILSDRFVEVPDMYGWTKSNVKTFAKWTGIDISFKGTDSGRVMKQSVDVGKSLKKIKKMTITLGD*

>PBP2x.32

MKKWQKYVLDYVVRDRRTPVENRVRVGQNMMLLTIFIFFIFIINFMIIIGTDQKFGVSLSEGAKKVYQETVTIQAKRGTIYDRNGTAIAVDSTTYSIYAILDKSFVSASDEKLYVQPSQYETVAYILKKHLGMKKTDVIKQLKRKGLFQVSFGPSGSGISYSTMSTIQKAMEDAKIKGIAFTTSPGRMYPNGTFASEFIGLASLTEDKKTGVKSLVGKTGLEASFDKILSGQDGVITYQKDRNGTTLLGTGKTVKKAIDGKDIYTTLSEPIQTFLETQMDVFQAKSNGQLASATLVNAKTGEILATTQRPTYNADTLKGLENTNYKWYSALHQGNFEPGSTMKVMTLAAAIDDKVFNPNETFSNANGLTIADATIQDWSINEGISTGQYMNYAQGFAFSSNVGMTKLEQKMGNAKWMNYLTKFRFGFPTRFGLKDEDAGIFPSDNIVTQAMSAFGQGISVTQIQMLRAFTAISNNGEMLEPQFISQIYDPNTASFRTANKEIVGKPVSKKAASETRQYMIGVGTDPEFGTLYSKTFGPIIKVGDLPVAVKSGTAQIGSEDGSGYQDGGLTNYVYSVVAMVPADKPDFLMYVTMTKPQHFGPLFWQDVVNPVLEEAYLMQDTLTKPVVSDANRQTTYKLPNFVGKNPGETSSELRRNLVQPVVLGTGSKIKKVSHQPGQTLTENQQVLILSDRFVEVPDMYGWTKSNVKTFAKWTGIDISFKGTDSGRVMKQSVDVGKSLKKIKKMTITLGD*

>PBP2x.53

MKKWQKYVLDYVVRDRRTPVENRVRVGQNMMLLTIFIFFIFIINFMIIIGTDQKFGVSLSEGAKKVYQETVTIQAKRGTIYDRNGTAIAVDSTTYSIYAILDKSFVSASDEKLYVQPSQYETVADILKKHLGMKKIDVIKQLKRKGLFQVSFGPSGSGISYSTMSTIQKAMEDAKIKGIAFTTSPGRMYPNGTFASEFIGLASLTEDKKTGVKSLVGKTGLEASFDKILSGQDGVITYQKDRNGTTLLGTGKTVKKAIDGKDIYTTLSEPIQTFLETQMDVFQAKSNGQLASATLVNAKTGEILATTQRPTYNADTLKGLENTNYKWYSALHQGNFEPGSTMKVMTLAAAIDDKVFNPNETFSNANGLTIADATIQDWSINEGISTGQYMNYAQGFAFSSNVGMTKLEQKMGNAKWMNYLTKFRFGFPTRFGLKDEDAGIFPSDNIVTQAMSAFGQGISVTQIQMLRAFTAISNNGEMLEPQFISQIYDPNTASFRTANKEIVGKPVSKKAASETRQYMIGVGTDPEFGTLYSKTFGPIIKVGDLPVAVKSGTAQIGSEDGSGYQDGGLTNYVYSVVAMVPADKPDFLMYVTMTKPQHFGPLFWQDVVNPVLEEAYLMQDTLTKPVVSDANRQTTYKLPNFVGKNPGETSSELRRNLVQPVVLGTGSKIKKVSHQPGQTLTENQQVLILSDRFVEVPDMYGWTKSNVKTFAKWTGIDISFKGTDSGRVMKQSVDVGKSLKKIKKMTITLGD*

>PBP2x.49

MKKWQKYVLDYVVRDRRTPVENRVRVGQNMMLLTIFIFFIFIINFMIIIGTDQKFGVSLSEGAKKVYQETVTIQAKRGTIYDRNGTAIAVDSTTYSIYAILDKSFVSASDEKLYVQPSQYETVADILKKHLGMKKTDVTKQLKRKGLFQVSFGPSGSGISYSTMSTIQKAMEDAKIKGIAFTTSPGRMYPNGTFASEFIGLASLTEDKKTGVKSLVGKTGLEASFDKILSGQDGVITYQKDRNGTTLLGTGKTVKKAIDGKDIYTTLSEPIQTFLETQMDVFQAKSNGQLASATLVNAKTGEILATTQRPTYNADTLKGLENTNYKWYSALHQGNFEPGSTMKVMTLAAAIDDKVFNPNETFSNANGLTIADATIQDWSINEGISTGQYMNYAQGFAFSSNVGMTKLEQKMGNAKWMNYLTKFRFGFPTRFGLKDEDAGIFPSDNIVTQAMSAFGQGISVTQIQMLRAFTAISNNGEMLEPQFISQIYDPNTASFRTANKEIVGKPVSKKAASETRQYMIGVGTDPEFGTLYSKTFGPIIKVGDLPVAVKSGTAQIGSEDGSGYQDGGLTNYVYSVVAMVPADKPDFLMYVTMTKPQHFGPLFWQDVVNPVLEEAYLMQDTLTKPVVSDANRQTTYKLPNFVGKNPGETSSELRRNLVQPVVLGTGSKIKKVSHQPGQTLTENQQVLILSDRFVEVPDMYGWTKSNVKTFAKWTGIDISFKGTDSGRVMKQSVDVGKSLKKIKKMTITLGD*

>PBP2x.48

MKKWQKYVLDYVVRDRRTPVENRVRVGQNMMLLTIFIFFIFIINFMIIIGTDQKFGVSLSEGAKKVYQETVTIQAKRGTIYDRNGTAIAVDSTTYSIYAILDKSFVSASDEKLYVQPSQYETVADILKKHLGMKKTDVIKQLKRKGLFQVSFGPSGSGISYSTMSTIQKAMEGAKIKGIAFTTSPGRMYPNGTFASEFIGLASLTEDKKTGVKSLVGKTGLEASFDKILSGQDGVITYQKDRNGTTLLGTGKTVKKAIDGKDIYTTLSEPIQTFLETQMDVFQAKSNGQLASATLVNAKTGEILATTQRPTYNADTLKGLENTNYKWYSALHQGNFEPGSTMKVMTLAAAIDDKVFNPNETFSNANGLTIADATIQDWSINEGISTGQYMNYAQGFAFSSNVGMTKLEQKMGNAKWMNYLTKFRFGFPTRFGLKDEDAGIFPSDNIVTQAMSAFGQGISVTQIQMLRAFTAISNNGEMLEPQFISQIYDPNTASFRTANKEIVGKPVSKKAASETRQYMIGVGTDPEFGTLYSKTFGPIIKVGDLPVAVKSGTAQIGSEDGSGYQDGGLTNYVYSVVAMVPADKPDFLMYVTMTKPQHFGPLFWQDVVNPVLEEAYLMQDTLTKPVVSDANRQTTYKLPNFVGKNPGETSSELRRNLVQPVVLGTGSKIKKVSHQPGQTLTENQQVLILSDRFVEVPDMYGWTKSNVKTFAKWTGIDISFKGTDSGRVMKQSVDVGKSLKKIKKMTITLGD*

>PBP2x.30

MKKWQKYVLDYVVRDRRTPVENRVRVGQNMMLLTIFIFFIFIINFMIIIGTDQKFGVSLSEGAKKVYQETVTIQAKRGTIYDRNGTAIAVDSTTYSIYAILDKSFVSASDEKLYVQPSQYETVADILKKHLGMKKTDVIKQLKRKGLFQVSFGPSGSGISYSTMSTIQKAMEDAKIKGIAFTTSPGRMYPNGTFASEFIGLASLTEDKKTGVKSLVGKTGLEASFNKILSGQDGVITYQKDRNGTTLLGTGKTVKKAIDGKDIYTTLSEPIQTFLETQMDVFQAKSNGQLASATLVNAKTGEILATTQRPTYNADTLKGLENTNYKWYSALHQGNFEPGSTMKVMTLAAAIDDKVFNPNETFSNANGLTIADATIQDWSINEGISTGQYMNYAQGFAFSSNVGMTKLEQKMGNAKWMNYLTKFRFGFPTRFGLKDEDAGIFPSDNIVTQAMSAFGQGISVTQIQMLRAFTAISNNGEMLEPQFISQIYDPNTASFRTANKEIVGKPVSKKAASETRQYMIGVGTDPEFGTLYSKTFGPIIKVGDLPVAVKSGTAQIGSEDGSGYQDGGLTNYVYSVVAMVPADKPDFLMYVTMTKPQHFGPLFWQDVVNPVLEEAYLMQDTLTKPVVSDANRQTTYKLPNFVGKNPGETSSELRRNLVQPVVLGTGSKIKKVSHQPGQTLTENQQVLILSDRFVEVPDMYGWTKSNVKTFAKWTGIDISFKGTDSGRVMKQSVDVGKSLKKIKKMTITLGD*

>PBP2x.51

MKKWQKYVLDYVVRDRRTPVENRVRVGQNMMLLTIFIFFIFIINFMIIIGTDQKFGVSLSEGAKKVYQETVTIQAKRGTIYDRNGTAIAVDSTTYSIYAILDKSFVSASDEKLYVQPSQYETVADILKKHLGMKKTDVIKQLKRKGLFQVSFGPSGSGISYSTMSTIQKAMEDAKIKGIAFTTSPGRMYPNGTFASEFIGLASLTEDKKTGVKSLVGKTGLEASFDKILSGQDGIITYQKDRNGTTLLGTGKTVKKAIDGKDIYTTLSEPIQTFLETQMDVFQAKSNGQLASATLVNAKTGEILATTQRPTYNADTLKGLENTNYKWYSALHQGNFEPGSTMKVMTLAAAIDDKVFNPNETFSNANGLTIADATIQDWSINEGISTGQYMNYAQGFAFSSNVGMTKLEQKMGNAKWMNYLTKFRFGFPTRFGLKDEDAGIFPSDNIVTQAMSAFGQGISVTQIQMLRAFTAISNNGEMLEPQFISQIYDPNTASFRTANKEIVGKPVSKKAASETRQYMIGVGTDPEFGTLYSKTFGPIIKVGDLPVAVKSGTAQIGSEDGSGYQDGGLTNYVYSVVAMVPADKPDFLMYVTMTKPQHFGPLFWQDVVNPVLEEAYLMQDTLTKPVVSDANRQTTYKLPNFVGKNPGETSSELRRNLVQPVVLGTGSKIKKVSHQPGQTLTENQQVLILSDRFVEVPDMYGWTKSNVKTFAKWTGIDISFKGTDSGRVMKQSVDVGKSLKKIKKMTITLGD*

>PBP2x.4

MKKWQKYVLDYVVRDRRTPVENRVRVGQNMMLLTIFIFFIFIINFMIIIGTDQKFGVSLSEGAKKVYQETVTIQAKRGTIYDRNGTAIAVDSTTYSIYAILDKSFVSASDEKLYVQPSQYETVADILKKHLGMKKTDVIKQLKRKGLFQVSFGPSGSGISYSTMSTIQKAMEDAKIKGIAFTTSPGRMYPNGTFASEFIGLASLTEDKKTGVKSLVGKTGLEASFDKILSGQDGVITYQKDRNGATLLGTGKTVKKAIDGKDIYTTLSEPIQTFLETQMDVFQAKSNGQLASATLVNAKTGEILATTQRPTYNADTLKGLENTNYKWYSALHQGNFEPGSTMKVMTLAAAIDDKVFNPNETFSNANGLTIADATIQDWSINEGISTGQYMNYAQGFAFSSNVGMTKLEQKMGNAKWMNYLTKFRFGFPTRFGLKDEDAGIFPSDNIVTQAMSAFGQGISVTQIQMLRAFTAISNNGEMLEPQFISQIYDPNTASFRTANKEIVGKPVSKKAASETRQYMIGVGTDPEFGTLYSKTFGPIIKVGDLPVAVKSGTAQIGSEDGSGYQDGGLTNYVYSVVAMVPADKPDFLMYVTMTKPQHFGPLFWQDVVNPVLEEAYLMQDTLTKPVVSDANRQTTYKLPNFVGKNPGETSSELRRNLVQPVVLGTGSKIKKVSHQPGQTLTENQQVLILSDRFVEVPDMYGWTKSNVKTFAKWTGIDISFKGTDSGRVMKQSVDVGKSLKKIKKMTITLGD*

>PBP2x.50

MKKWQKYVLDYVVRDRRTPVENRVRVGQNMMLLTIFIFFIFIINFMIIIGTDQKFGVSLSEGAKKVYQETVTIQAKRGTIYDRNGTAIAVDSTTYSIYAILDKSFVSASDEKLYVQPSQYETVADILKKHLGMKKTDVIKQLKRKGLFQVSFGPSGSGISYSTMSTIQKAMEDAKIKGIAFTTSPGRMYPNGTFASEFIGLASLTEDKKTGVKSLVGKTGLEASFDKILSGQDGVITYQKDRNGATLLGTGKTVKKAIDGKDIYTTLSEPIQTFLETQMDVFQAKSNGQLASATLVNAKTGEILATTQRPTYNADTLKGLENTNYKWYSALHQGNFEPGSTMKVMTLAAAIDDKVFNPNETFSNANGLTIADATIQDWSINEGISTGQYMNYAQGFAFSSNVGMTKLEQKMGNAKWMNYLTKFRFGFPTRFGLKDEDAGIFPSDNIVTQAMSAFGQGISVTQIQMLRAFTAISNNGEMLEPQFISQIYDPNTASFRTANKEIVGKPVSKKAASETRQYMISVGTDPEFGTLYSKTFGPIIKVGDLPVAVKSGTAQIGSEDGSGYQDGGLTNYVYSVVAMVPADKPDFLMYVTMTKPQHFGPLFWQDVVNPVLEEAYLMQDTLTKPVVSDANRQTTYKLPNFVGKNPGETSSELRRNLVQPVVLGTGSKIKKVSHQPGQTLTENQQVLILSDRFVEVPDMYGWTKSNVKTFAKWTGIDISFKGTDSGRVMKQSVDVGKSLKKIKKMTITLGD*

>PBP2x.29

MKKWQKYVLDYVVRDRRTPVENRVRVGQNMMLLTIFIFFIFIINFMIIIGTDQKFGVSLSEGAKKVYQETVTIQAKRGTIYDRNGTAIAVDSTTYSIYAILDKSFVSASDEKLYVQPSQYETVADILKKHLGMKKTDVIKQLKRKGLFQVSFGPSGSGISYSTMSTIQKAMEDAKIKGIAFTTSPGRMYPNGTFASEFIGLASLTEDKKTGVKSLVGKTGLEASFDKILSGQDGVITYQKDRNGATLLGTGKTVKKAIDGKDIYTTLSEPIQTFLETQMDVFQAKSNGQLASATLVNAKTGEILATTQRPTYNADTLKGLENTNYKWYSALHQGNFEPGSTMKVMTLAAAIDDKVFNPNETFSNANGLTIADATIQDWSINEGISTGQYMNYAQGFAFSSNVGMTKLEQKMGNAKWMNYLTKFRFGFPTRFGLKDEDAGIFPSDNIVTQAMSAFGQGISVTQIQMLRAFTAISNNGEMLEPQFISQIYDPNTASFRTANKEIVGKPVSKKAASETRQYMIGVGTDPEFGTLYSKTFGPIIKVGDLPVAVKSGTAQIGSEDGSGYQDGGLTNYVYSVVAMVPADKPDFLMYVTMTKPQHFGPLFWQDVVNPVLEEAYLMQDTLTKPVVSDANRQTAYKLPNFVGKNPGETSSELRRNLVQPVVLGTGSKIKKVSHQPGQTLTENQQVLILSDRFVEVPDMYGWTKSNVKTFAKWTGIDISFKGTDSGRVMKQSVDVGKSLKKIKKMTITLGD*

>PBP2x.86

MKKWQKYVLDYVVRDRRTPVENRVRVGQNMMLLTIFIFFIFIINFMIIIGTDQKFGVSLSEGAKKVYQETVTIQAKRGTIYDRNGTAIAVDSTTYSIYAILDKSFVSASDEKLYVQPSQYETVADILKKHLGMKKTDVIKQLKRKGLFQVSFGPSGSGISYSTMSTIQKAMEDAKIKGIAFTTSPGRMYPNGTFASEFIGLASLTEDKKTGVKSLVGKTGLEASFDKILSGQDGVITYQKDRNGTTLLGTGKTVKKAIDGKDIYTTLSDPIQTFLETQMDVFQAKSNGQLASATLVNAKTGEILATTQRPTYNADTLKGLENTNYKWYSALHQGNFEPGSTMKVMTLAAAIDDKVFNPNETFSNANGLTIADATIQDWSINEGISTGQYMNYAQGFAFSSNVGMTKLEQKMGNAKWMNYLTKFRFGFPTRFGLKDEDAGIFPSDNIVTQAMSAFGQGISVTQIQMLRAFTAISNNGEMLEPQFISQIYDPNTASFRTANKEIVGKPVSKKAASETRQYMIGVGTDPEFGTLYSKTFGPIIKVGDLPVAVKSGTAQIGSEDGSGYQDGGLTNYVYSVVAMVPADKPDFLMYVTMTKPQHFGPLFWQDVVNPVLEEAYLMQDTLTKPVVSDANRQTTYKLPNFVGKNPGETSSELRRNLVQPVVLGTGSKIKKVSHQPGQTLTENQQVLILSDRFVEVPDMYGWTKSNVKTFAKWTGIDISFKGTDSGRVMKQSVDVGKSLKKIKKMTITLGD*

>PBP2x.93

MKKWQKYVLDYVVRDRRTPVENRVRVGQNMMLLTIFIFFIFIINFMIIIGTDQKFGVSLSEGAKKVYQETVTIQAKRGTIYDRNGTAIAVDSTTYSIYAILDKSFVSASDEKLYVQPSQYETVADILKKHLGMKKTDVIKQLKRKGLFQVSFGPSGSGISYSTMSTIQKAMEDAKIKGIAFTTSPGRMYPNGTFASEFIGLASLTEDKKTGVKSLVGKTGLEASFDKILSGQDGVITYQKDRNGTTLLGTGKTVKKAIDGKDIYTTLSEPIQTFLETQMDVFQAKSNGQLASAALVNAKTGEILATTQRPTYNADTLKGLENTNYKWYSALHQGNFEPGSTMKVMTLAAAIDAKVFNPNETFSNANGLTIADATIQDWSINEGISTGQYMNYAQGFAFSSNVGMTKLEQKMGNAKWMNYLTKFRFGFPTRFGLKDEDAGIFPSDNIVTQAMSAFGQGISVTQIQMLRAFTAISNNGEMLEPQFISQIYDPNTASFRTANKEIVGKPVSKKAASETRQYMIGVGTDPEFGTLYSKTFGPIIKVGDLPVAVKSGTAQIGSEDGSGYQDGGLTNYVYSVVAMVPADKPDFLMYVTMTKPQHFGPLFWQDVVNPVLEEAYLMQDTLTKPVVSDANRQTTYKLPNFVGKNPGETSSELRRNLVQPVVLGTGSKIKKVSHQSGQTLTENQQVLILSDRFVEVPDMYGWTKSNVETFAKWTGIDISFKGTDSGRVMKQSVDVGKSLKKIKKMTITLGD*

>PBP2x.76

MKKWQKYVLDYVVRDRRTPVENRVRVGQNMMLLTIFIFFIFIINFMIIIGTDQKFGVSLSEGAKKVYQETVTIQAKRGTIYDRNGTAIAVDSTTYSIYAILDKSFVSASDEKLYVQPSQYETVADILKKHLGMKKTDVIKQLKRKGLFQVSFGPSGSGISYSTMSTIQKAMEDAKIKGIAFTTSPGRMYPNGTFASEFIGLASLTEDKKTGVKSLVGKTGLEASFDKILSGQDGVITYQKDRNGTTLLGTGKTVKKAIDGKDIYTTLSEPIQTFLETQMDVFQAKSNGQLASATLVNAKTGEILATAQRPTYNADTLKGLENTNYKWYSALHQGNFEPGSTMKVMTLAAAIDDKVFNPNETFSNANGLTIADATIQDWSINEGISTGQYMNYAQGFAFSSNVGMTKLEQKMGNAKWMNYLTKFRFGFPTRFGLKDEDAGIFPSDNIVTQAMSAFGQGISVTQIQMLRAFTAISNNGEMLEPQFISQIYDPNTASFRTANKEIVGKPVSKKAASETRQYMIGVGTDPEFGTLYSKTFGPIIKVGDLPVAVKSGTAQIGSEDGSGYQDGGLTNYVYSVVAMVPADKPDFLMYVTMTKPQHFGPLFWQDVVNPVLEEAYLMQDTLTKPVVSDANRQTTYKLPNFVGKNPGETSSELRRNLVQPVVLGTGSKIKKVSHQPGQTLTENQQVLILSDRFVEVPDMYGWTKSNVKTFAKWTGIDISFKGTDSGRVMKQSVDVGKSLKKIKKMTITLGD*

>PBP2x.81

MKKWQKYVLDYVVRDRRTPVENRVRVGQNMMLLTIFIFFIFIINFMIIIGTDQKFGVSLSEGAKKVYQETVTIQAKRGTIYDRNGTAIAVDSTTYSIYAILDKSFVSASDEKLYVQPSQYETVADILKKHLGMKKTDVIKQLKRKGLFQVSFGPSGSGISYSTMSTIQKAMEDAKIKGIAFTTSPGRMYPNGTFASEFIGLASLTEDKKTGVKSLVGKTGLEASFDKILSGQDGVITYQKDRNGTTLLGTGKTVKKAIDGKDIYTTLSEPIQTFLETQMDVFQAKSNGQLASATLVNAKTGEILATTQRPTYNADTLKGLENTNYKWYSTLHQGNFEPGSTMKVMTLAAAIDDKVFNPNETFSNANGLTIADATIQDWSINEGISTGQYMNYAQGFAFSSNVGMTKLEQKMGNAKWMNYLTKFRFGFPTRFGLKDEDAGIFPSDNIVTQAMSAFGQGISVTQIQMLRAFTAISNNGEMLEPQFISQIYDPNTASFRTANKEIVGKPVSKKAASETRQYMIGVGTDPEFGTLYSKTFGPIIKVGDLPVAVKSGTAQIGSEDGSGYQDGGLTNYVYSVVAMVPADKPDFLMYVTMTKPQHFGPLFWQDVVNPVLEEAYLMQDTLTKPVVSDANRQTTYKLPNFVGKNPGETSSELRRNLVQPVVLGTGSKIKKVSHQPGQTLTENQQVLILSDRFVEVPDMYGWTKSNVKTFAKWTGIDISFKGTDSGRVMKQSVDVGKSLKKIKKMTITLGD*

>PBP2x.42

MKKWQKYVLDYVVRDRRTPVENRVRVGQNMMLLTIFIFFIFIINFMIIIGTDQKFGVSLSEGAKKVYQETVTIQAKRGTIYDRNGTAIAVDSTTYSIYAILDKSFVSASDEKLYVQPSQYETVADILKKHLGMKKTDVIKQLKRKGLFQVSFGPSGSGISYSTMSTIQKAMEDAKIKGIAFTTSPGRMYPNGTFASEFIGLASLTEDKKTGVKSLVGKTGLEASFDKILSGQDGVITYQKDRNGTTLLGTGKTVKKAIDGKDIYTTLSEPIQTFLETQMDVFQAKSNGQLASATLVNAKTGEILATTQRPTYNADTLKGLENTNYKWYSALHQGNFEPGSTMKVMTLAAAIDAKVFNPNETFSNANGLTIADATIQDWSINEGISTGQYMNYAQGFAFSSNVGMTKLEQKMGNAKWMNYLTKFRFGFPTRFGLKDEDAGIFPSDNIVTQAMSAFGQGISVTQIQMLRAFTAISNNGEMLEPQFISQIYDPNTASFRTANKEIVGKPVSKKAASETRQYMIGVGTDPEFGTLYSKTFGPIIKVGDLPVAVKSGTAQIGSEDGSGYQDGGLTNYVYSVVAMVPADKPDFLMYVTMTKPQHFGPLFWQDVVNPVLEEAYLMQDTLTKPVVSDANRQTTYKLPNFVGKNPGETSSELRRNLVQPVVLGTGSKIKKVSHQPGQTLTENQQVLILSDRFVEVPDMYGWTKSNVKTFAKWTGIDISFKGTDSGRVMKQSVDVGKSLKKIKKMTITLGD*

>PBP2x.94

MKKWQKYVLDYVVRDRRTPVENRVRVGQNMMLLTIFIFFIFIINFMIIIGTDQKFGVSLSEGAKKVYQETVTIQAKRGTIYDRNGTAIAVDSTTYSIYAILDKSFVSASDEKLYVQPSQYETVADILKKHLGMKKTDVIKQLKRKGLFQVSFGPSGSGISYSTMSTIQKAMEDAKIKGIAFTTSPGRMYPNGTFASEFIGLASLTEDKKTGVKSLVGKTGLEASFDKILSGQDGVITYQKDRNGTTLLGTGKTVKKAIDGKDIYTTLSEPIQTFLETQMDVFQAKSNGQLASATLVNAKTGEILATTQRPTYNADTLKGLENTNYKWYSALHQGNFEPGSTMKVMTLAAAIDDKVFNPNETFSNANGLTIADATIQDWSINEGISTGQYMNYAQGFVFSSNVGMTKLEQKMGNAKWMNYLTKFRFGFPTRFGLKDEDAGIFPSDNIVTQAMSAFGQGISVTQIQMLRAFTAISNNGEMLEPQFISQIYDPNTASFRTANKEVVGKPVSKKAASETRQYMIGVGTDPEFGTLYSKTFGPIIKVGDLPVAVKSGTAQIGSEDGSGYQDGGLTNYVYSVVAMVPADKPDFLMYVTMTKPQHFGPLFWQDVVNPVLEEAYLMQDTLTKPVVSDANRQTTYKLPNFVGKNPGETSSELRRNLVQPVVLGTGSKIKKVSHQPGQTLTENQQVLILSDRFVEVPDMYGWTKSNVKTFAKWTGIDISFKGTDSGRVMKQSVDVGKSLKKIKKMTITLGD*

>PBP2x.106

MKKWQKYVLDYVVRDRRTPVENRVRVGQNMMLLTIFIFFIFIINFMIIIGTDQKFGVSLSEGAKKVYQETVTIQAKRGTIYDRNGTAIAVDSTTYSIYAILDKSFVSASDEKLYVQPSQYETVADILKKHLGMKKTDVIKQLKRKGLFQVSFGPSGSGISYSTMSTIQKAMEDAKIKGIAFTTSPGRMYPNGTFASEFIGLASLTEDKKTGVKSLVGKTGLEASFDKILSGQDGVITYQKDRNGTTLLGTGKTVKKAIDGKDIYTTLSEPIQTFLETQMDVFQAKSNGQLASATLVNAKTGEILATTQRPTYNADTLKGLENTNYKWYSALHQGNFEPGSTMKVMTLAAAIDDKVFNPNETFSNANGLTIADATIQDWSINEGISTGQYMNYAQGFAFSSNVGMTKLEQKMGNAKWMNYLTKFRFGFPTRFGLKDEDAGIFPSDNIVTQAMSAFGQGISVTQIQMLRAFTAISNNGEMLEPQFISQIYDPNTASFRTANKEVVGKPVSKKAASETRQYMIGVGTDPEFGTLYSKTFGPIIKVGDLPVAVKSGTAQIGSEDGSGYQDGGLTNYVYSVVAMVPADKPDFLMYVTMTKPQHFGPLFWQDVVNPVLEEAYLMQDTLTKPVVSDANRQTTYKLPNFVGKNPGETSSELRRNLVQPVVLGTGSKIKKVSHQPGQTLTENQQVLILSDRFVEVPDMYGWTKSNVKTFAKWTGIDISFKGTDSGRVMKQSVDVGKSLKKIKKMTITLGD*

>PBP2x.3

MKKWQKYVLDYVVRDRRTPVENRVRVGQNMMLLTIFIFFIFIINFMIIIGTDQKFGVSLSEGAKKVYQETVTIQAKRGTIYDRNGTAIAVDSTTYSIYAILDKSFVSASDEKLYVQPSQYETVADILKKHLGMKKTDVIKQLKRKGLFQVSFGPSGSGISYSTMSTIQKAMEDAKIKGIAFTTSPGRMYPNGTFASEFIGLASLTEDKKTGVKSLVGKTGLEASFDKILSGQDGVITYQKDRNGTTLLGTGKTVKKAIDGKDIYTTLSEPIQTFLETQMDVFQAKSNGQLASATLVNAKTGEILATTQRPTYNADTLKGLENTNYKWYSALHQGNFEPGSTMKVMTLAAAIDDKVFNPNETFSNANGLTIADATIQDWSINEGISTGQYMNYAQGFAFSSNVGMTKLEQKMGNAKWMNYLTKFRFGFPTRFGLKDEDAGIFPSDNIVTQAMSAFGQGISVTQIQMLRAFTAISNNGEMLEPQFISQIYDPNTASFRTANKEVVGKPVSKKAASETRQYMIGVGTDPEFGTLYSKTFGPIIKVGDLPVAVKSGTAQIGSEDGSGYQDGGLTNYVYSVVAMVPADKPDFLMYVTMTKPQHFGPLFWQDVVNPVLEEAYLMQDTLTKPVVSDANRQTTYKLPNFVGKNPGETSSELRRNLVQPVVLGTGSKIKKVSHQSGQTLTENQQVLILSDRFVEVPDMYGWTKSNVETFAKWTGIDISFKGTDSGRVMKQSVDVGKSLKKIKKMTITLGD*

>PBP2x.31

MKKWQKYVLDYVVRDRRTPVENRVRVGQNMMLLTIFIFFIFIINFMIIIGTDQKFGVSLSEGAKKVYQETVTIQAKRGTIYDRNGTAIAVDSTTYSIYAILDKSFVSASDEKLYVQPSQYETVADILKKHLGMKKTDVIKQLKRKGLFQVSFGPSGSGISYSTMSTIQKAMEDAKIKGIAFTTSPGRMYPNGTFASEFIGLASLTEDKKTGVKSLVGKTGLEASFDKILSGQDGVITYQKDRNGTTLLGTGKTVKKAIDGKDIYTTLSEPIQTFLETQMDVFQAKSNGQLASATLVNAKTGEILATTQRPTYNADTLKGLENTNYKWYSALHQGNFEPGSTMKVMTLAAAIDDKVFNPNETFSNANGLTIADATIQDWSINEGISTGQYMNYAQGFAFSSNVGMTKLEQKMGNAKWMNYLTKFRFGFPTRFGLKDEDAGIFPSDNIVTQAMSAFGQGISVTQIQMLRAFTAISNNGEMLEPQFISQIYDPNTASFRTANKEVVGKPVSKKAASETRQYMIGVGTDPEFGTLYSKTFGPIIKVGDLPVAVKSGTAQIGSEDGSGYQDGGLTNYVYSVVAMVPADKPDFLMYVTMTKPQHFGPLFWQDVVNPVLEEAYLMQDTLTKPVVSDANRQTTYKLPNFVGKNPGETSSELRRNLVQPVVLGTGSKIKKVSHQSGQTLTENQQVLILSDRFVEVPDMYGWTKSNVETFAKWTGIDISFKGTDSGRVIKQSVDVGKSLKKIKKMTITLGD*

>PBP2x.78

MKKWQKYVLDYVVRDRRTPVENRVRVGQNMMLLTIFIFFIFIINFMIIIGTDQKFGVSLSEGAKKVYQETVTIQAKRGTIYDRNGTAIAVDSTTYSIYAILDKSFVSASDEKLYVQPSQYETVADILKKHLGMKKTDVIKQLKRKGLFQVSFGPSGSGISYSTMSTIQKAMEDAKIKGIAFTTSPGRMYPNGTFASEFIGLASLTEDKKTGVKSLVGKTGLEASFDKILSGQDGVITYQKDRNGTTLLGTGKTVKKAIDGKDIYTTLSEPIQTFLETQMDVFQAKSNGQLASATLVNAKTGEILATTQRPTYNADTLKGLENTNYKWYSALHQGNFEPGSTMKVMTLAAAIDDKVFNPNETFSNANGLTIADATIQDWSINEGISTGQYMNYAQGFAFSSNVGMTKLEQKMGNAKWMNYLTKFRFGFPTRFGLKDEDAGIFPSDNIVTQAMSAFGQGISVTQIQMLRAFTAISNNGEMLEPQFISQIYDPNTASFRTANKEIIGKPVSKKAASETRQYMIGVGTDPEFGTLYSKTFGPIIKVGDLPVAVKSGTAQIGSEDGSGYQDGGLTNYVYSVVAMVPADKPDFLMYVTMTKPQHFGPLFWQDVVNPVLEEAYLMQDTLTKPVVSDANRQTTYKLPNFVGKNPGETSSELRRNLVQPVVLGTGSKIKKVSHQPGQTLTENQQVLILSDRFVEVPDMYGWTKSNVKTFAKWTGIDISFKGTDSGRVMKQSVDVGKSLKKIKKMTITLGD*

>PBP2x.85

MKKWQKYVLDYVVRDRRTPVENRVRVGQNMMLLTIFIFFIFIINFMIIIGTDQKFGVSLSEGAKKVYQETVTIQAKRGTIYDRNGTAIAVDSTTYSIYAILDKSFVSASDEKLYVQPSQYETVADILKKHLGMKKTDVIKQLKRKGLFQVSFGPSGSGISYSTMSTIQKAMEDAKIKGIAFTTSPGRMYPNGTFASEFIGLASLTEDKKTGVKSLVGKTGLEASFDKILSGQDGVITYQKDRNGTTLLGTGKTVKKAIDGKDIYTTLSEPIQTFLETQMDVFQAKSNGQLASATLVNAKTGEILATTQRPTYNADTLKGLENTNYKWYSALHQGNFEPGSTMKVMTLAAAIDDKVFNPNETFSNANGLTIADATIQDWSINEGISTGQYMNYAQGFAFSSNVGMTKLEQKMGNAKWMNYLTKFRFGFPTRFGLKDEDAGIFPSDNIVTQAMSAFGQGISVTQIQMLRAFTAISNNGEMLEPQFISQIYDPNTASFRTANKEIVGKPVSKKAASETRQYMIGVGTDSEFGTLYSKTFGPIIKVGDLPVAVKSGTAQIGSEDGSGYQDGGLTNYVYSVVAMVPADKPDFLMYVTMTKPQHFGPLFWQDVVNPVLEEAYLMQDTLTKPVVSDANRQTTYKLPNFVGKNPGETSSELRRNLVQPVVLGTGSKIKKVSHQPGQTLTENQQVLILSDRFVEVPDMYGWTKSNVKTFAKWTGIDISFKGTDSGRVMKQSVDVGKSLKKIKKMTITLGD*

>PBP2x.103

MKKWQKYVLDYVVRDRRTPVENRVRVGQNMMLLTIFIFFIFIINFMIIIGTDQKFGVSLSEGAKKVYQETVTIQAKRGTIYDRNGTAIAVDSTTYSIYAILDKSFVSASDEKLYVQPSQYETVADILKKHLGMKKTDVIKQLKRKGLFQVSFGPSGSGISYSTMSTIQKAMEDAKIKGIAFTTSPGRMYPNGTFASEFIGLASLTEDKKTGVKSLVGKTGLEASFDKILSGQDGVITYQKDRNGTTLLGTGKTVKKAIDGKDIYTTLSEPIQTFLETQMDVFQAKSNGQLASATLVNAKTGEILATTQRPTYNADTLKGLENTNYKWYSALHQGNFEPGSTMKVMTLAAAIDDKVFNPNETFSNANGLTIADATIQDWSINEGISTGQYMNYAQGFAFSSNVGMTKLEQKMGNAKWMNYLTKFRFGFPTRFGLKDEDAGIFPSDNIVTQAMSAFGQGISVTQIQMLRAFTAISNNGEMLEPQFISQIYDPNTASFRTANKEIVGKPVSKKAASETRQYMIGVGTDPEFGTLYSKIFGPIIKVGDLPVAVKSGTAQIGSEDGSGYQDGGLTNYVYSVVAMVPADKPDFLMYVTMTKPQHFGPLFWQDVVNPVLEEAYLMQDTLTKPVVSDANRQTTYKLPNFVGKNPGETSSELRRNLVQPVVLGTGSKIKKVSHQPGQTLTENQQVLILSDRFVEVPDMYGWTKSNVKTFAKWTGIDISFKGTDSGRVMKQSVDVGKPLKKIKKMTITLGD*

>PBP2x.12

MKKWQKYVLDYVVRDRRTPVENRVRVGQNMMLLTIFIFFIFIINFMIIIGTDQKFGVSLSEGAKKVYQETVTIQAKRGTIYDRNGTAIAVDSTTYSIYAILDKSFVSASDEKLYVQPSQYETVADILKKHLGMKKTDVIKQLKRKGLFQVSFGPSGSGISYSTMSTIQKAMEDAKIKGIAFTTSPGRMYPNGTFASEFIGLASLTEDKKTGVKSLVGKTGLEASFDKILSGQDGVITYQKDRNGTTLLGTGKTVKKAIDGKDIYTTLSEPIQTFLETQMDVFQAKSNGQLASATLVNAKTGEILATTQRPTYNADTLKGLENTNYKWYSALHQGNFEPGSTMKVMTLAAAIDDKVFNPNETFSNANGLTIADATIQDWSINEGISTGQYMNYAQGFAFSSNVGMTKLEQKMGNAKWMNYLTKFRFGFPTRFGLKDEDAGIFPSDNIVTQAMSAFGQGISVTQIQMLRAFTAISNNGEMLEPQFISQIYDPNTASFRTANKEIVGKPVSKKAASETRQYMIGVGTDPEFGTLYSKTFGPIIKVGDLPVAVKSGTAQIGSEDGTGYQDGGLTNYVYSVVAMVPADKPDFLMYVTMTKPQHFGPLFWQDVVNPVLEEAYLMQDTLTKPVVSDANRQTTYKLPNFVGKNPGETSSELRRNLVQPVVLGTGSKIKKVSHQPGQTLTENQQVLILSDRFVEVPDMYGWTKSNVKTFAKWTGIDISFKGTDSGRVMKQSVDVGKSLKKIKKMTITLGD*

>PBP2x.28

MKKWQKYVLDYVVRDRRTPVENRVRVGQNMMLLTIFIFFIFIINFMIIIGTDQKFGVSLSEGAKKVYQETVTIQAKRGTIYDRNGTAIAVDSTTYSIYAILDKSFVSASDEKLYVQPSQYETVADILKKHLGMKKTDVIKQLKRKGLFQVSFGPSGSGISYSTMSTIQKAMEDAKIKGIAFTTSPGRMYPNGTFASEFIGLASLTEDKKTGVKSLVGKTGLEASFDKILSGQDGVITYQKDRNGTTLLGTGKTVKKAIDGKDIYTTLSEPIQTFLETQMDVFQAKSNGQLASATLVNAKTGEILATTQRPTYNADTLKGLENTNYKWYSALHQGNFEPGSTMKVMTLAAAIDDKVFNPNETFSNANGLTIADATIQDWSINEGISTGQYMNYAQGFAFSSNVGMTKLEQKMGNAKWMNYLTKFRFGFPTRFGLKDEDAGIFPSDNIVTQAMSAFGQGISVTQIQMLRAFTAISNNGEMLEPQFISQIYDPNTASFRTANKEIVGKPVSKKAASETRQYMIGVGTDPEFGTLYSKTFGPIIKVGDLPVAVKSGTAQIGSEDGTGYQDGGLTNYVYSVVAMVPADKPDFLMYVTMTKPQHFGPLFWQDVVNPVLEEAYLMQDTLTKPVVSDANRQTTYKLPNFVGKNPGETSSELRRNLVQPVVLGTGSKIKKVSHQSGQTLTENQQVLILSDRFVEVPDMYGWTKSNVETFAKWTGIDISFKGTDSGRVMKQSVDVGKSLKKIKKMTITLGD*

>PBP2x.63

MKKWQKYVLDYVVRDRRTPVENRVRVGQNMMLLTIFIFFIFIINFMIIIGTDQKFGVSLSEGAKKVYQETVTIQAKRGTIYDRNGTAIAVDSTTYSIYAILDKSFVSASDEKLYVQPSQYETVADILKKHLGMKKTDVIKQLKRKGLFQVSFGPSGSGISYSTMSTIQKAMEDAKIKGIAFTTSPGRMYPNGTFASEFIGLASLTEDKKTGVKSLVGKTGLEASFDKILSGQDGVITYQKDRNGTTLLGTGKTVKKAIDGKDIYTTLSEPIQTFLETQMDVFQAKSNGQLASATLVNAKTGEILATTQRPTYNADTLKGLENTNYKWYSALHQGNFEPGSTMKVMTLAAAIDDKVFNPNETFSNANGLTIADATIQDWSINEGISTGQYMNYAQGFAFSSNVGMTKLEQKMGNAKWMNYLTKFRFGFPTRFGLKDEDAGIFPSDNIVTQAMSAFGQGISVTQIQMLRAFTAISNNGEMLEPQFISQIYDPNTASFRTANKEIVGKPVSKKAASETRQYMIGVGTDPEFGTLYSKTFGPIIKVGDLPVAVKSGTAQIGSEDGSGYQDGGLTNYVYSVVAMVPADKPDFLMYVTTTKPQHFGPLFWQDVVNPVLEEAYLMQDTLTKPVVSDANRQTTYKLPNFVGKNPGETSSELRRNLVQPVVLGTGSKIKKVSHQPGQTLTENQQVLILSDRFVEVPDMYGWTKSNVKTFAKWTGIDISFKGTDSGRVMKQSVDVGKSLKKIKKMTITLGD*

>PBP2x.54

MKKWQKYVLDYVVRDRRTPVENRVRVGQNMMLLTIFIFFIFIINFMIIIGTDQKFGVSLSEGAKKVYQETVTIQAKRGTIYDRNGTAIAVDSTTYSIYAILDKSFVSASDEKLYVQPSQYETVADILKKHLGMKKTDVIKQLKRKGLFQVSFGPSGSGISYSTMSTIQKAMEDAKIKGIAFTTSPGRMYPNGTFASEFIGLASLTEDKKTGVKSLVGKTGLEASFDKILSGQDGVITYQKDRNGTTLLGTGKTVKKAIDGKDIYTTLSEPIQTFLETQMDVFQAKSNGQLASATLVNAKTGEILATTQRPTYNADTLKGLENTNYKWYSALHQGNFEPGSTMKVMTLAAAIDDKVFNPNETFSNANGLTIADATIQDWSINEGISTGQYMNYAQGFAFSSNVGMTKLEQKMGNAKWMNYLTKFRFGFPTRFGLKDEDAGIFPSDNIVTQAMSAFGQGISVTQIQMLRAFTAISNNGEMLEPQFISQIYDPNTASFRTANKEIVGKPVSKKAASETRQYMIGVGTDPEFGTLYSKTFGPIIKVGDLPVAVKSGTAQIGSEDGSGYQDGGLTNYVYSVVAMVPADKPDFLMYVTMTKPQHFDPLFWQDVVNPVLEEAYLMQDTLTKPVVSDANRQTTYKLPNFVGKNPGETSSELRRNLVQPVVLGTGSKIKKVSHQPGQTLTENQQVLILSDRFVEVPDMYGWTKSNVKTFAKWTGIDISFKGTDSGRVMKQSVDVGKSLKKIKKMTITLGD*

>PBP2x.44

MKKWQKYVLDYVVRDRRTPVENRVRVGQNMMLLTIFIFFIFIINFMIIIGTDQKFGVSLSEGAKKVYQETVTIQAKRGTIYDRNGTAIAVDSTTYSIYAILDKSFVSASDEKLYVQPSQYETVADILKKHLGMKKTDVIKQLKRKGLFQVSFGPSGSGISYSTMSTIQKAMEDAKIKGIAFTTSPGRMYPNGTFASEFIGLASLTEDKKTGVKSLVGKTGLEASFDKILSGQDGVITYQKDRNGTTLLGTGKTVKKAIDGKDIYTTLSEPIQTFLETQMDVFQAKSNGQLASATLVNAKTGEILATTQRPTYNADTLKGLENTNYKWYSALHQGNFEPGSTMKVMTLAAAIDDKVFNPNETFSNANGLTIADATIQDWSINEGISTGQYMNYAQGFAFSSNVGMTKLEQKMGNAKWMNYLTKFRFGFPTRFGLKDEDAGIFPSDNIVTQAMSAFGQGISVTQIQMLRAFTAISNNGEMLEPQFISQIYDPNTASFRTANKEIVGKPVSKKAASETRQYMIGVGTDPEFGTLYSKTFGPIIKVGDLPVAVKSGTAQIGSEDGSGYQDGGLTNYVYSVVAMVPADKPDFLMYVTMTKPQHFAHLFWQDVVNPVLEEAYLMQDTLTKPVVSDANRQTTYKLPNFVGKNPGETSSELRRNLVQPVVLGTGSKIKKVSHQPGQTLTENQQVLILSDRFVEVPDMYGWTKSNVKTFAKWTGIDISFKGTDSGRVMKQSVDVGKSLKKIKKMTITLGD*

>PBP2x.22

MKKWQKYVLDYVVRDRRTPVENRVRVGQNMMLLTIFIFFIFIINFMIIIGTDQKFGVSLSEGAKKVYQETVTIQAKRGTIYDRNGTAIAVDSTTYSIYAILDKSFVSASDEKLYVQPSQYETVADILKKHLGMKKTDVIKQLKRKGLFQVSFGPSGSGISYSTMSTIQKAMEDAKIKGIAFTTSPGRMYPNGTFASEFIGLASLTEDKKTGVKSLVGKTGLEASFDKILSGQDGVITYQKDRNGTTLLGTGKTVKKAIDGKDIYTTLSEPIQTFLETQMDVFQAKSNGQLASATLVNAKTGEILATTQRPTYNADTLKGLENTNYKWYSALHQGNFEPGSTMKVMTLAAAIDDKVFNPNETFSNANGLTIADATIQDWSINEGISTGQYMNYAQGFAFSSNVGMTKLEQKMGNAKWMNYLTKFRFGFPTRFGLKDEDAGIFPSDNIVTQAMSAFGQGISVTQIQMLRAFTAISNNGEMLEPQFISQIYDPNTASFRTANKEIVGKPVSKKAASETRQYMIGVGTDPEFGTLYSKTFGPIIKVGDLPVAVKSGTAQIGSEDGSGYQDGGLTNYVYSVVAMVPADKPDFLMYVTMTKPQHFGLLFWQDVVNPVLEEAYLMQDTLTKPVVSDANRQTTYKLPNFVGKNPGETSSELRRNLVQPVVLGTGSKIKKVSHQPGQTLTENQQVLILSDRFVEVPDMYGWTKSNVKTFAKWTGIDISFKGTDSGRVMKQSVDVGKSLKKIKKMTITLGD*

>PBP2x.98

MKKWQKYVLDYVVRDRRTPVENRVRVGQNMMLLTIFIFFIFIINFMIIIGTDQKFGVSLSEGAKKVYQETVTIQAKRGTIYDRNGTAIAVDSTTYSIYAILDKSFVSASDEKLYVQPSQYETVADILKKHLGMKKTDVIKQLKRKGLFQVSFGPSGSGISYSTMSTIQKAMEDAKIKGIAFTTSPGRMYPNGTFASEFIGLASLTEDKKTGVKSLVGKTGLEASFDKILSGQDGVITYQKDRNGTTLLGTGKTVKKAIDGKDIYTTLSEPIQTFLETQMDVFQAKSNGQLASATLVNAKTGEILATTQRPTYNADTLKGLENTNYKWYSALHQGNFEPGSTMKVMTLAAAIDDKVFNPNETFSNANGLTIADATIQDWSINEGISTGQYMNYAQGFAFSSNVGMTKLEQKMGNAKWMNYLTKFRFGFPTRFGLKDEDAGIFPSDNIVTQAMSAFGQGISVTQIQMLRAFTAISNNGEMLEPQFISQIYDPNTASFRTANKEIVGKPVSKKAASETRQYMIGVGTDPEFGTLYSKTFGPIIKVGDLPVAVKSGTAQIGSEDGSGYQDGGLTNYVYSVVAMVPADKPDFLMYVTMTKPQHFGPLFWQDVVNPVLEEAYLMQDTLTKPVVSDANHQTTYKLPNFVGKNPGETSSELRRNLVQPVVLGTGSKIKKVSHQPGQTLTENQQVLILSDRFVEVPDMYGWTKSNVKTFAKWTGIDISFKGTDSGRVMKQSVDVGKSLKKIKKMTITLGD*

>PBP2x.83

MKKWQKYVLDYVVRDRRTPVENRVRVGQNMMLLTIFIFFIFIINFMIIIGTDQKFGVSLSEGAKKVYQETVTIQAKRGTIYDRNGTAIAVDSTTYSIYAILDKSFVSASDEKLYVQPSQYETVADILKKHLGMKKTDVIKQLKRKGLFQVSFGPSGSGISYSTMSTIQKAMEDAKIKGIAFTTSPGRMYPNGTFASEFIGLASLTEDKKTGVKSLVGKTGLEASFDKILSGQDGVITYQKDRNGTTLLGTGKTVKKAIDGKDIYTTLSEPIQTFLETQMDVFQAKSNGQLASATLVNAKTGEILATTQRPTYNADTLKGLENTNYKWYSALHQGNFEPGSTMKVMTLAAAIDDKVFNPNETFSNANGLTIADATIQDWSINEGISTGQYMNYAQGFAFSSNVGMTKLEQKMGNAKWMNYLTKFRFGFPTRFGLKDEDAGIFPSDNIVTQAMSAFGQGISVTQIQMLRAFTAISNNGEMLEPQFISQIYDPNTASFRTANKEIVGKPVSKKAASETRQYMIGVGTDPEFGTLYSKTFGPIIKVGDLPVAVKSGTAQIGSEDGSGYQDGGLTNYVYSVVAMVPADKPDFLMYVTMTKPQHFGPLFWQDVVNPVLEEAYLMQDTLTKPVVSDANRQTTYKLPNFVGKNPGETSSELRRNLVQPVVLGTGSKIKKVSHQSGQTLTENQQVLILSDRFVEVPDMYGWTKSNVETFAKWTGIDISFKGTDSGRVMKQSVDVGKSLKKIKKMTITLGD*

>PBP2x.99

MKKWQKYVLDYVVRDRRTPVENRVRVGQNMMLLTIFIFFIFIINFMIIIGTDQKFGVSLSEGAKKVYQETVTIQAKRGTIYDRNGTAIAVDSTTYSIYAILDKSFVSASDEKLYVQPSQYETVADILKKHLGMKKTDVIKQLKRKGLFQVSFGPSGSGISYSTMSTIQKAMEDAKIKGIAFTTSPGRMYPNGTFASEFIGLASLTEDKKTGVKSLVGKTGLEASFDKILSGQDGVITYQKDRNGTTLLGTGKTVKKAIDGKDIYTTLSEPIQTFLETQMDVFQAKSNGQLASATLVNAKTGEILATTQRPTYNADTLKGLENTNYKWYSALHQGNFEPGSTMKVMTLAAAIDDKVFNPNETFSNANGLTIADATIQDWSINEGISTGQYMNYAQGFAFSSNVGMTKLEQKMGNAKWMNYLTKFRFGFPTRFGLKDEDAGIFPSDNIVTQAMSAFGQGISVTQIQMLRAFTAISNNGEMLEPQFISQIYDPNTASFRTANKEIVGKPVSKKAASETRQYMIGVGTDPEFGTLYSKTFGPIIKVGDLPVAVKSGTAQIGSEDGSGYQDGGLTNYVYSVVAMVPADKPDFLMYVTMTKPQHFGPLFWQDVVNPVLEEAYLMQDTLTKPVVSDANRQTTYKLPNFVGKNPGETSSELRRNLVQPVVLGTGSKIKKVSHQPGQTLTENQQVLILSDRFVEIPDMYGWTKSNVKTFAKWTGIDISFKGTDSGRVMKQSVDVGKSLKKIKKMTITLGD*

>PBP2x.46

MKKWQKYVLDYVVRDRRTPVENRVRVGQNMMLLTIFIFFIFIINFMIIIGTDQKFGVSLSEGAKKVYQETVTIQAKRGTIYDRNGTAIAVDSTTYSIYAILDKSFVSASDEKLYVQPSQYETVADILKKHLGMKKTDVIKQLKRKGLFQVSFGPSGSGISYSTMSTIQKAMEDAKIKGIAFTTSPGRMYPNGTFASEFIGLASLTEDKKTGVKSLVGKTGLEASFDKILSGQDGVITYQKDRNGTTLLGTGKTVKKAIDGKDIYTTLSEPIQTFLETQMDVFQAKSNGQLASATLVNAKTGEILATTQRPTYNADTLKGLENTNYKWYSALHQGNFEPGSTMKVMTLAAAIDDKVFNPNETFSNANGLTIADATIQDWSINEGISTGQYMNYAQGFAFSSNVGMTKLEQKMGNAKWMNYLTKFRFGFPTRFGLKDEDAGIFPSDNIVTQAMSAFGQGISVTQIQMLRAFTAISNNGEMLEPQFISQIYDPNTASFRTANKEIVGKPVSKKAASETRQYMIGVGTDPEFGTLYSKTFGPIIKVGDLPVAVKSGTAQIGSEDGSGYQDGGLTNYVYSVVAMVPADKPDFLMYVTMTKPQHFGPLFWQDVVNPVLEEAYLMQDTLTKPVVSDANRQTTYKLPNFVGKNPGETSSELRRNLVQPVVLGTGSKIKKVSHQPGQTLTENQQVLILSDRFVEVPDMYSWTKSNVKTFAKWTGIDISFKGTDSGRVMKQSVDVGKSLKKIKKMTITLGD*

>PBP2x.101

MKKWQKYVLDYVVRDRRTPVENRVRVGQNMMLLTIFIFFIFIINFMIIIGTDQKFGVSLSEGAKKVYQETVTIQAKRGTIYDRNGTAIAVDSTTYSIYAILDKSFVSASDEKLYVQPSQYETVADILKKHLGMKKTDVIKQLKRKGLFQVSFGPSGSGISYSTMSTIQKAMEDAKIKGIAFTTSPGRMYPNGTFASEFIGLASLTEDKKTGVKSLVGKTGLEASFDKILSGQDGVITYQKDRNGTTLLGTGKTVKKAIDGKDIYTTLSEPIQTFLETQMDVFQAKSNGQLASATLVNAKTGEILATTQRPTYNADTLKGLENTNYKWYSALHQGNFEPGSTMKVMTLAAAIDDKVFNPNETFSNANGLTIADATIQDWSINEGISTGQYMNYAQGFAFSSNVGMTKLEQKMGNAKWMNYLTKFRFGFPTRFGLKDEDAGIFPSDNIVTQAMSAFGQGISVTQIQMLRAFTAISNNGEMLEPQFISQIYDPNTASFRTANKEIVGKPVSKKAASETRQYMIGVGTDPEFGTLYSKTFGPIIKVGDLPVAVKSGTAQIGSEDGSGYQDGGLTNYVYSVVAMVPADKPDFLMYVTMTKPQHFGPLFWQDVVNPVLEEAYLMQDTLTKPVVSDANRQTTYKLPNFVGKNPGETSSELRRNLVQPVVLGTGSKIKKVSHQPGQTLTENQQVLILSDRFVEVPDMYGWTKSNVKTFAKWTGIDISFKGTDSGRVMKQSVDIGKSLKKIKKMTITLGD*
